# Biophysical characterization of calcium-binding and modulatory-domain dynamics in a pentameric ligand-gated ion channel

**DOI:** 10.1101/2022.05.06.490775

**Authors:** Marie Lycksell, Urška Rovšnik, Anton Hanke, Anne Martel, Rebecca J Howard, Erik Lindahl

**Affiliations:** Department of Biochemistry and Biophysics, Science for Life Laboratory, Stockholm University, Stockholm, Sweden; Institute of Pharmacy and Molecular Biotechnology (IPMB), Heidelberg University, Heidelberg 69120, Germany; Institut Laue-Langevin, Grenoble, France; Department of Applied Physics, Science for Life Laboratory, KTH Royal Institute of Technology, Stockholm, Sweden

## Abstract

Pentameric ligand-gated ion channels (pLGICs) perform electrochemical signal transduction in organisms ranging from bacteria to humans. Among the prokaryotic pLGICs there is architectural diversity involving N-terminal domains (NTDs) not found in eukaryotic relatives, exemplified by the calcium-sensitive channel (DeCLIC) from a *Desulfofustis* deltaproteobacterium, which has an NTD in addition to the canonical pLGIC structure. Here we have characterized the structure and dynamics of DeCLIC through cryo-electron microscopy (cryo-EM), small-angle neutron scattering (SANS), and molecular dynamics (MD) simulations. In the presence and absence of calcium, cryo-EM yielded structures with alternative conformations of the calcium binding site. SANS profiles further revealed conformational diversity at room temperature beyond that observed in static structures, shown through MD to be largely attributable to rigid body motions of the NTD relative to the protein core, with expanded and asymmetric conformations improving the fit of the SANS data. This work reveals the range of motion available to the DeCLIC NTD and calcium binding site, expanding the conformational landscape of the pLGIC family. Further, these findings demonstrate the power of combining low-resolution scattering, high-resolution structural, and MD-simulation data to elucidate interfacial interactions that are highly conserved in the pLGIC family.

## Introduction

Pentameric ligand-gated ion channels (pLGICs) belong to a family of membrane proteins responsible for fast signal transduction, utilized by numerous organisms from bacteria to humans [1]. In response to a chemical stimulus, these proteins undergo a conformational change, allowing ion flux across the cell membrane [2]. This state-dependent ion permeation is especially important for the proper function of the mammalian nervous system; accordingly, dysfunction in these channels is linked to a variety of neurodegenerative disorders such as epilepsy, hyperekplexia, Alzheimer’s and Parkinson’s diseases [3]. These effects make pLGICs critical targets for a number of therapeutic agents, including anesthetics, benzodiazepines and neurosteroids [4, 5]. A detailed understanding of channel pharmacology and conformational change depends on high-quality structural data, potentially complemented by dynamic information from methods like molecular dynamics (MD) simulations and small-angle scattering [6].

In recent years, advances in X-ray crystallography and cryo-electron microscopy (cryo-EM) [7], have revealed several conserved features of pLGIC structure [8]. All pLGICs share a five-fold homo- or heteromeric assembly. Each subunit contains a transmembrane domain (TMD) consisting of four helices (M1–M4), and an extracellular domain (ECD) consisting of ten beta strands (β1–β10) and a characteristic Pro-(prokaryotes) or Cys-loop (eukaryotes) [9]. Most eukaryotic pLGICs also contain an intracellular domain (ICD) of variable length [2]. In contrast, several prokaryotic family members contain an additional amino-terminal domain (NTD) [1, 9] not observed in any eukaryotic homologs. Despite these insights, the details of ion permeation and modulation in this family, and the structure-function relationships of these peripheral domains, remain poorly understood.

Similar to several eukaryotic ion channels [10], the bacterial channel DeCLIC — derived from a Desulfofustis deltaproteobacterium – was recently shown to be modulated by circulating calcium ions [11]. DeCLIC was crystallized in both the presence and absence of calcium, offering models in apparent closed and open states (Figure 1A). When crystallized in high (150 mM) calcium, DeCLIC bound calcium via several electronegative groups in the ECD periphery, and the permeation pore in this structure was too narrow to conduct ions at multiple points along the TMD (Figure 1A-B). In the absence of calcium, the pore instead adopted a wide-open *apo* state, notably distinct from related pLGIC structures (Figure 1A-B). The ECD vestibule was also tightly constricted, raising questions about the pathway of ion transit, and about the general relevance of these structures in the gating cycle of DeCLIC or other family members. Notably, both structures included a partially resolved NTD, to the best of our knowledge the first one determined in any pLGIC. Although not seen in eukaryotic pLGICs, N-terminal domains occur in other ligand-gated channels such as AMPA and NMDA receptors [12], where they can bind ligands and have allosteric effects [13]. In the case of DeCLIC, the NTD was found to influence gating kinetics, with each subunit containing two jelly-roll lobes (NTD1 and NTD2) packed against the peripheral ECD (Figure 1A). However, details about functional and structural relevance of this domain remain unclear [11].

**Figure 1:**
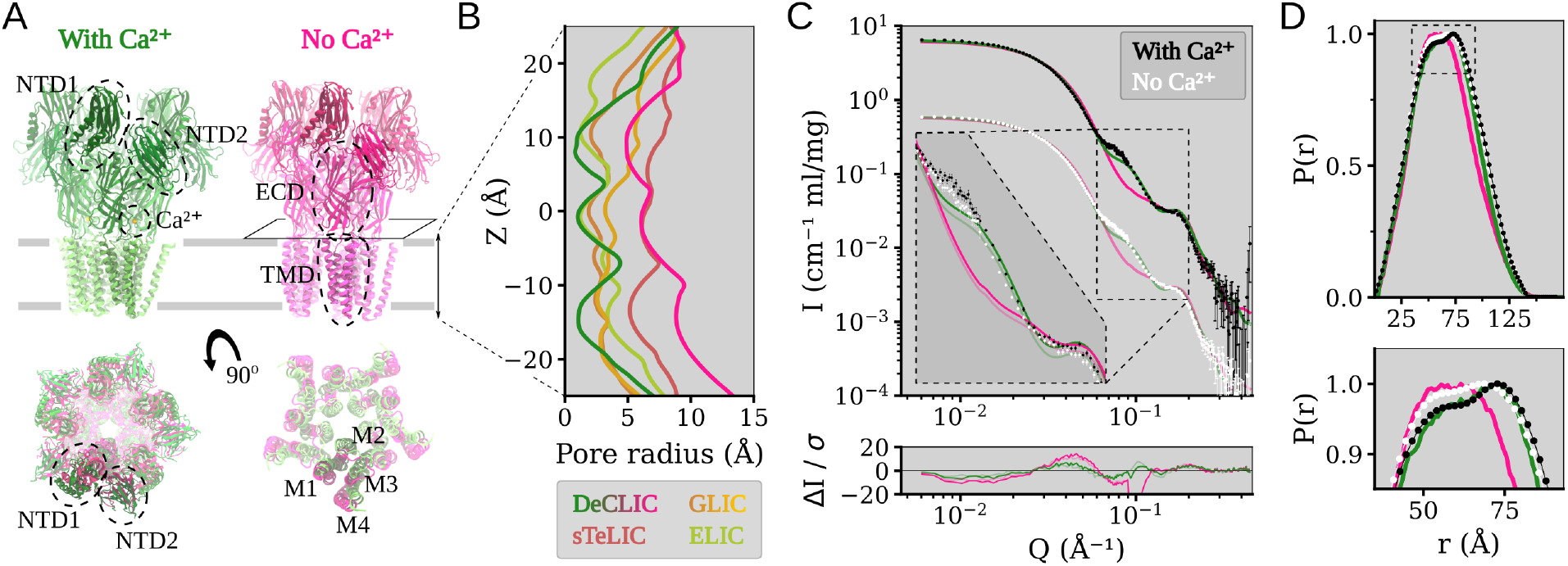
Structure overview of DeCLIC, and the fit of the DeCLIC X-ray structures to small angle neutron scattering data. (A) The structure of DeCLIC consists of two beta-rich N-terminal domains (NTD1 and NTD2), a betasandwich extracellular domain (ECD) with a calcium binding site, and a transmembrane domain (TMD) with four helices M1 through M4. X-ray crystallography structures have been determined by Hu et al. [11] in the presence (green) and absence (pink) of calcium, with absence of calcium leading to a conformation with a more contracted NTD and a more expanded TMD. (B) Pore profiles of the TMD region in DeCLIC (green and pink), and of the related channels GLIC (brown and yellow), sTeLIC (brick), and ELIC (lime). In the presence of calcium, DeCLIC (green) has a contracted pore similar to ELIC (lime) and GLIC at resting conditions (brown). In the absence of calcium, DeCLIC (pink) has a wide pore at the 9’ hydrophobic gate (Z=0 Å), similar to sTeLIC (brick), and is at the 16’ contraction (Z≈10 Å) similar to GLIC at activating conditions (yellow). (C) Small angle neutron scattering from DeCLIC in the presence (black) and absence (white) of calcium, with predicted curves from the crystal structures in the presence (green) and absence (pink) of calcium. The data-set in the presence of calcium has been offset by a factor of 10 for clarity. The inset shows the data on the same scale zoomed in on *Q*∈ [0.06, 0.2] Å ^-1^, with darker curves fitted to the SANS data with calcium, and paler curves fitted to the calcium-free SANS data. The calcium-free crystal structure has a poor goodness of fit (χ^2^ > 60) to the SANS data, while the calcium-containing crystal structure give a moderate goodness of fit (χ^2^ of 10.8 to SANS with calcium, χ^2^ of 8.8 to SANS without calcium), deviating from the experimental data mainly for *Q*∈[0.06, 0.09] Å ^-1^. (D) Pair distance distribution of DeCLIC from SANS with (black) and without (white) calcium, and from crystal structures with (green) and without (pink) calcium. There is close agreement between the distribution from SANS with calcium and from the calcium-containing crystal structure, while the SANS data collected in the absence of calcium have features similar to both crystal structures (zoom box).

Here, we sought to understand the mechanisms of ion binding and permeation, the physiological relevance of resolved states, and the dynamic behavior of novel domains in DeCLIC, using three complementary biophysical methods. In particular, we applied small-angle neutron scattering (SANS) to obtain low-resolution information on the room temperature solution structure, single-particle cryo-EM to determine higher-resolution structural information that avoid crystal contacts, and molecular dynamics simulations for observing dynamics and conformational sampling on atomic level. Our data support a dominant population of closed channels in both the presence and absence of calcium, and a mechanism for calcium block via a peripheral site at the ECD-subunit interface. We further find the spontaneous translation of the NTD lobes, especially NTD1, helps describe some features of our SANS data and support a new detailed model of pLGIC conformational diversity.

## Results

### Small-angle neutron scattering reveals deviations from static structures

To investigate solution-phase structural properties of DeCLIC in the presence and absence of inhibitory calcium, we collected SANS data in either 10 mM CaCl_2_ (Ca^2+^) or 10 mM EDTA (*apo*) conditions. To optimize for homogeneous monodisperse samples, data were collected from a single peak from an inline size-exclusion column [6]. As calculated by Guinier analysis, the sample radius of gyration (R_g_) was slightly larger under Ca^2+^ conditions than *apo* conditions (52.0±0.2 Å compared to 51.6±0.2 Å). Similarly, R_g_ calculated from X-ray structures in the presence and absence of Ca^2+^ (49.5 Å and 48.2 Å, respectively) was also larger in the presence of Ca^2+^ (Supplementary Figure S1, Supplementary Tables S1, S2). Molecular weight estimates in Ca^2+^ and *apo* conditions (348 kD and 318 kD, respectively) were within 10% of that predicted from the amino acid sequence (342 kD), which is within the expected uncertainty arising from concentration determination. Accordingly, we proceeded to compare fits of the full scattering profiles to curves predicted from past X-ray structures.

Scattering profiles under Ca^2+^ and *apo* conditions were similar, and both were better described by the closed X-ray structure (χ^2^=10.8 for Ca^2+^, χ^2^=8.8 for *apo*) than by the open structure (χ^2^>60 for data collected under either condition) (Figure 1C). Deviation from the closed-state curve was primarily evident for a feature around *Q*∈ [0.06, 0.09] Å ^-1^ (Figure 1C, dashed inset). As real-space distances relate to momentum transfer by *d* = 2*π/*Q, this feature corresponds to a range of 70–105 Å, relatively long-range compared to the better-matched feature around *Q*∈ [0.10, 0.20] Å ^-1^. Overall, the distribution of pairwise distances was well described by the closed X-ray structure (Figure 1D), though it did show a slight underestimation of the prevalence of distances >100 Å (Supplementary Figure S2A) that could indicate the presence of more expanded conformations. The pairwise distance distribution from SANS data under *apo* conditions deviated modestly from both calcium-SANS data and the closed structure, with an increased contribution of shorter distances approaching the distribution function for the open structure. Thus, SANS curves under both Ca^2+^ and *apo* conditions were consistent with predominantly closed populations of channels, with some deviations at relatively long pairwise distances, and in the overall distribution of distances in the *apo* state.

### Differential closed cryo-EM structures in the presence and absence of calcium

To complement our SANS curves, we next collected cryo-EM data for DeCLIC in the presence and absence of 10 mM Ca^2+^, with the aim of sampling the channel’s conformational space without crystal contacts or highmM calcium. Each dataset yielded a single C5-symmetric reconstruction, with comparable resolution (3.5 Å and 3.2 Å in the presence and absence of calcium, respectively) (Figure 2A, Supplementary Figures S3, Fig S4, Fig S5, Table S3). Classification was also attempted without symmetry, but yielded only lower-resolution densities. For both conditions, the resolution was higher in the TMD and ECD, but worsened to *≥*4.5 Å in the NTD (Figure 2A). Whereas the backbone and majority of side chains were clearly resolved in the TMD and ECD (Supplementary Figure S6), some residues could not be built in NTD1 and the NTD1-NTD2 loop (Supplementary Table S4). The quality of the NTD was moderately better in the absence of calcium, with more residues resolved in both lobes (Supplementary Table S4).

**Figure 2:**
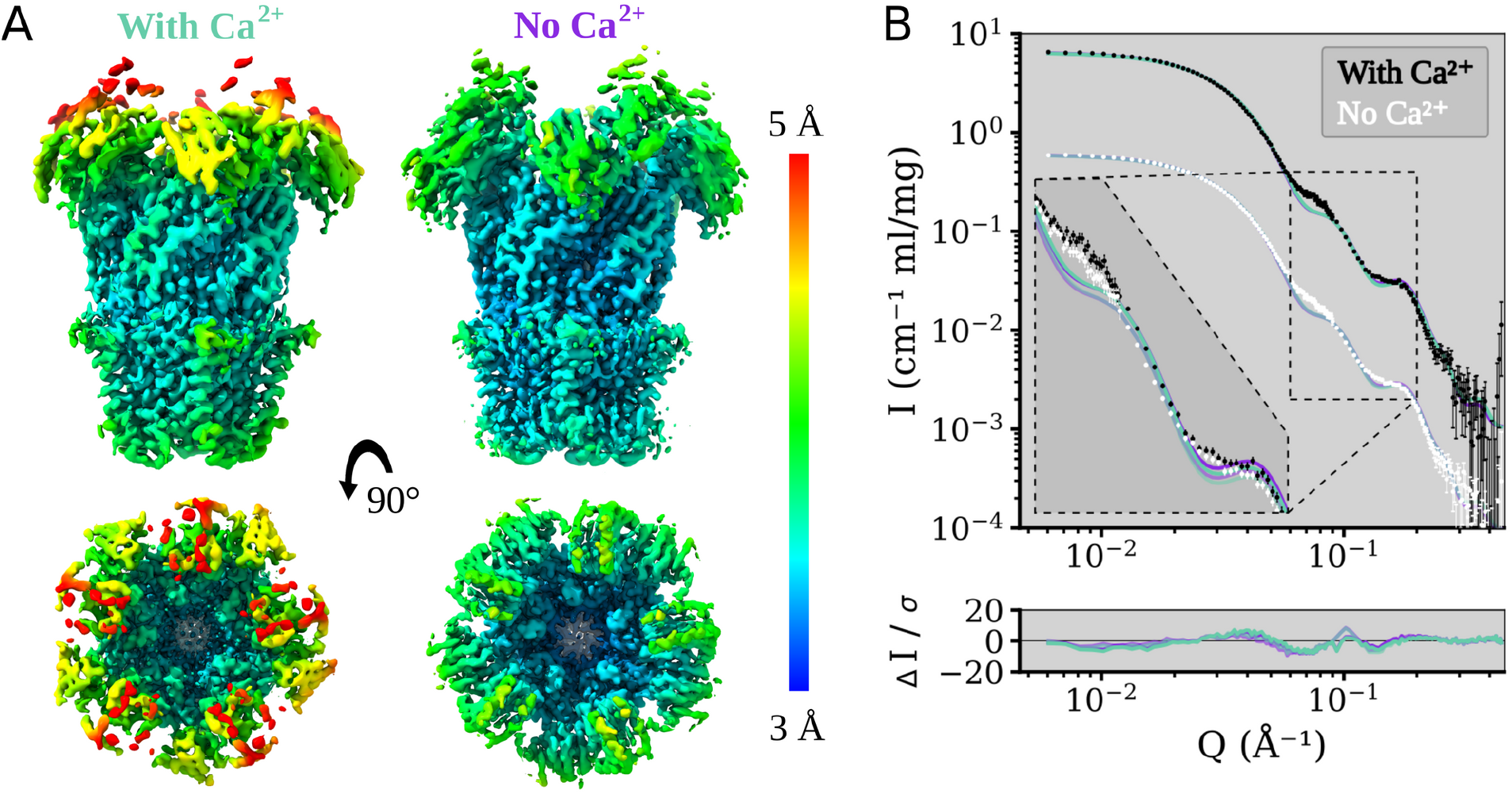
Local resolution colouring of the DeCLIC Cryo-EM reconstructions in the presence and absence of calcium and their agreement with the SANS data. (A) Cryo-EM density for DeCLIC with 10 mM Ca^2+^ present was resolved to an overall resolution of 3.5 Å (left) and density where Ca^2+^ was absent from the experimental conditions was resolved to 3.2 Å (right). Density is colored by local resolution according to scale bar to the right. The quality is worse in the NTD region of the protein compared to that of the TMD or ECD. (B) Fits of model spectra calculated from the two cryo-EM structures in the presence (aquamarine) and absence (purple) of Ca^2+^ to SANS data collected with Ca^2+^ (black), and with no Ca^2+^ (white). The structures yield similar goodness of fit, χ^2^∈ [11.2, 12.1]. SANS data with Ca^2+^ is offset by a factor of 10 for easier visualization. The inset shows the data on the same scale zoomed on *Q*∈ [0.06, 0.2] Å ^-1^, with darker curves fitted to the with-calcium SANS data, and paler curves fitted to the calcium-free SANS data.

Independent models were built manually in the two conditions. The greatest difference between them was around each ECD-subunit interface, particularly involving the β1-β2 loop and loop F, as detailed in the next section. Otherwise, the two models were largely superimposable (RMSD ≤ 0.63 Å) particularly in the TMD, with similarly closed pores. Indeed, the pore profiles of both models included hydrophobic constrictions to ≤1 Å radius at both L554 (9’) and F561 (16’) in the pore-lining M2 helices (Supplementary Figure S7A), consistent with features of previous X-ray structures [11]. The ECD vestibule in both models featured an additional constriction to *∼*5 Å radius, defined primarily by residue N405 in the β4–β5 loop (Supplementary Figure S7B). A weaker density in this vestibule could be attributed to an alternative, inward-facing, rotamer of residue W407. Interestingly, W407 was previously reported to orient into the ECD vestibule upon crystallization in the absence of Ca^2+^, leading to an even tighter constriction potentially incompatible with ion conduction via a linear ECD pathway [11].

Next, we compared theoretical curves predicted from our cryo-EM models to the SANS data described above (Fig 2B). Similar fits were obtained for curves based on either the calcium-bound (χ^2^=12.1 for Ca^2+^-SANS, χ^2^=11.3 for *apo*-SANS) or calcium-free cryo-EM models (χ^2^=11.2 for Ca^2+^, χ^2^=11.7 for *apo*). Pairwise distribution functions for the two cryo-EM models were also comparable (Supplementary Figure S2A). However, similar to the closed X-ray structure, a feature in the range *Q*∈ [0.06, 0.09] Å ^-1^ was poorly fit by both cryo-EM models (Figure 2B, *inset*). Thus, cryo-EM data broadly confirmed our SANS curves in substantiating a predominant population of closed channels, while leaving some features of the scattering data unexplained.

### Ion interactions at the calcium site revealed by cryo-EM and molecular dynamics

Although cryo-EM models of DeCLIC in the presence and absence of calcium were similar in apparent conductance states, they exhibited notable differences at each ECD-subunit interface. The quality of the densities in this region allowed building of the complete backbone and sidechains of the β1-β2 loop on the principal side of each interface, and of loop F on the complementary side (Figure 3A–B). The dataset with calcium also featured a small non-protein density between these loops (Figure 3B). Because this density was absent in the calcium-free dataset, and located near several acidic side chains (E347, E479, E480, E481), it was interpreted as a Ca^2+^ ion. In particular, carboxylate atoms of residues E347 and E480 (Figure 3B, Supplementary Figure S8B-C) were positioned <3 Å from the modeled ion, primed to coordinate it from both sides (Figure 3B, Figure S8B-C). In the calcium-free model, this ion density was absent, and E480 adopted a different rotamer facing away from the subunit interface (Figure 3A–B). Accordingly, the distance between E347 and E480 increased from <6 Å to >8 Å.

**Figure 3:**
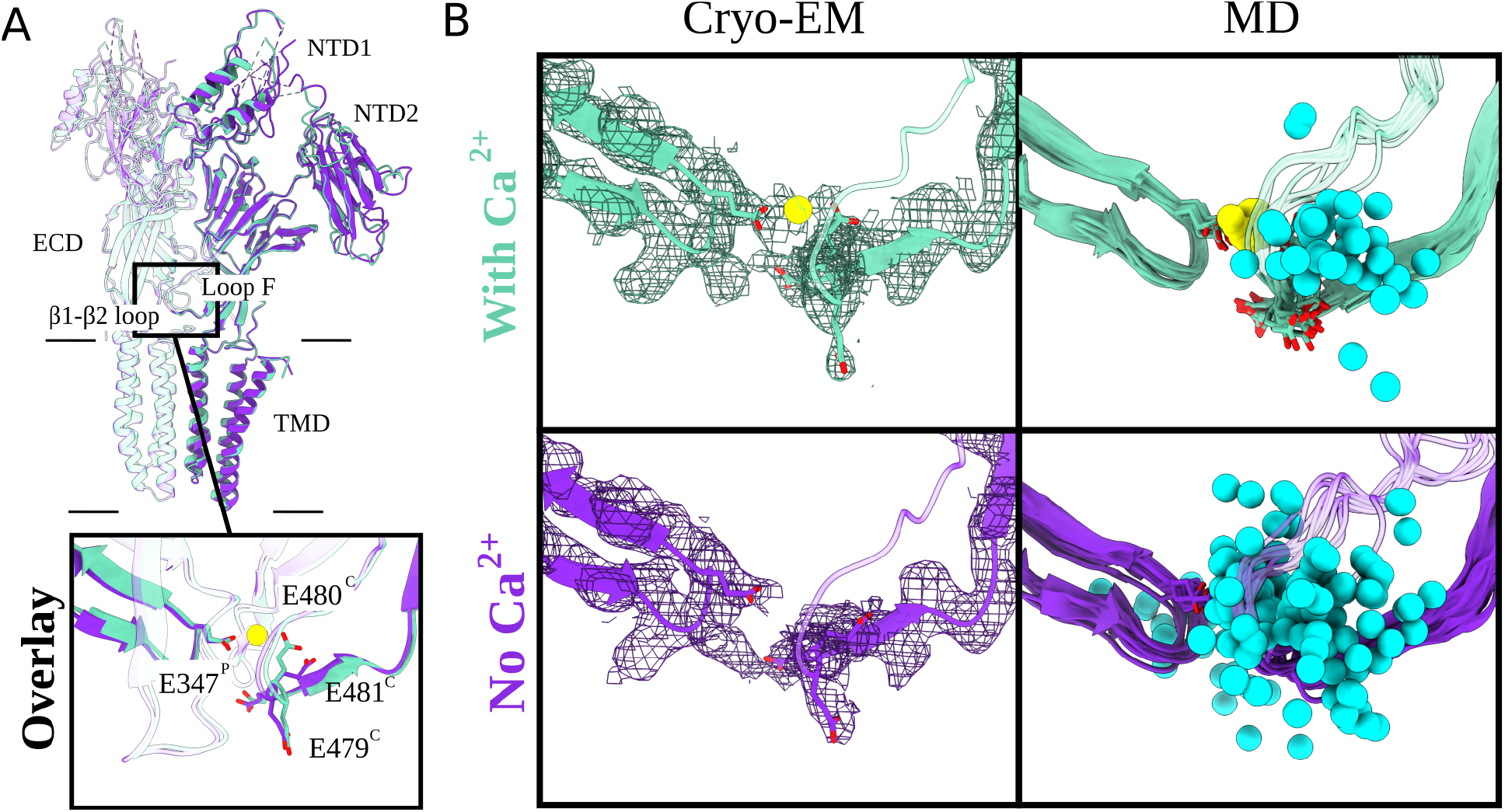
Arrangement of the calcium-binding residues elucidated by cryo-EM and MD. (A) Overlay of two adjacent subunits of DeCLIC cryo-EM structures with and without Ca^2+^ (aquamarine and purple, respectively). Protein domains and main Ca^2+^ binding motifs are shown. Those include the transmembrane domain (TMD), extracellular domain (ECD) and two amino terminal domains (NTD1, NTD2), as well as the β1–β2 loop from the principal subunit (P) and loop F from the complementary subunit (C). The boxed-in view (*bottom*) shows the relevant sidechains (sticks, colored by heteroatom). The Ca^2+^ ion present in the Ca^2+^ dataset (aquamarine) is colored in yellow. (B) Cryo-EM panels (*left*) represent the cryo-EM structures and their corresponding densities (mesh at σ = 0.012 - 0.015). Relevant sidechains around the Ca^2+^ binding site (loop F (C), β1–β2 loop (P)) are displayed as sticks (heteroatom coloring). The density for the Ca^2+^ binding residues as well as for the Ca^2+^ ion (in With Ca^2+^ condition) is well resolved. MD panels (*right*) illustrate eleven individual conformations (one snapshot for every 100 ns) from 1 *µ*s long MD simulations of the cryo-EM models. The relevant Ca^2+^ residues are displayed as sticks (colored by heteroatom). In cyan we have Na^+^ ions (sphere, trajectories generated for every 10 ns) and Ca^2+^ ion in yellow. In the no-Ca^2+^ condition the Na^+^ ions are free to enter the cavity which is occupied by Ca^2+^ in the with Ca^2+^ condition.

To confirm the stability of this putative calcium-binding site, we performed quadruplicate 1-*µ*s all-atom molecular dynamics simulations of each cryo-EM model, inserted in a lipid bilayer and solvated in water and sodium chloride. These simulations relaxed to stable conformational ensembles within 200 ns in the ECD and TMD (C*α* RMSD <2 Å, Supplementary Figure S9D–E); the NTD was substantially more variable, as described in subsequent sections. During simulations of the model with calcium, the distance between putative calcium-coordinating residues E347 and E480 remained <7 Å (Supplementary Figure S8B–C, *center*). Proximal residues E479 and E481 were also stably oriented within 13 Å of E347, possibly contributing to the negative electrostatic environment of the binding site (Supplementary Figure S8B–C, *left and right*). Moreover, all five calcium ions remained bound in all simulations, and no other ions were seen to transit the interface between subunits (Figure 3B). Conversely, in simulations of the cryo-EM model without calcium, E480 sampled various orientations up to 10 Å from E347 (Supplementary Figure S8B–C, *center*). Residues in loop F also fluctuated more in simulations of the calcium-free versus calcium-bound cryo-EM models (Supplementary Figure S10). Interestingly, the unoccupied calcium site at each subunit interface was frequently filled by sodium ions, in some cases transiting to or from the ECD vestibule (Figure 3B, Supplementary Movie S1).

To assess the possible influence of experimental conditions and models obtained with different techniques, we also ran simulations of the previously reported closed X-ray structure [11]. This structure was chosen due to its overall similarity to both our cryo-EM models, and was simulated in the presence and absence of calcium ions at the proposed binding site. In the X-ray structure, calcium was primarily coordinated between E347 and E479, rather than E480 (Supplementary Figure S8A). Nonetheless—echoing simulations of our cryo-EM models—the distance between coordinating residues remained below 6 Å with calcium present, and increased to more than 10 Å with calcium removed (Supplementary Figure S87B–C, *left*). Removing calcium from the X-ray structure also increased fluctuations in loop F (Supplementary Figure S10). Thus, structural and simulations data supported a stable binding site for calcium at the β1-β2/loop-F subunit interface, with unbinding mobilizing loop F and allowing transit of monovalent ions.

### Simulations demonstrate relative instability of open versus closed states

Simulations of closed DeCLIC structures determined by either cryo-EM or X-ray crystallography were notably stable in the TMD (Figure 4A, G; Supplementary Figure S7A), with only minor deviations in the peripheral M4 helices (Figure 4C; Supplementary Figure S11). Although no open state was evident in our cryo-EM data, we also sought to compare dynamics of open DeCLIC by simulating a previously reported X-ray structure with a wide-open pore (Hu et al., 2020). However, all simulations of this system exhibited substantial asymmetric deviations in TMD architecture, largely associated with the sizable gaps located between subunits in the initial structure (Figure 4B, D). Placing lipids into a partially resolved upper-leaflet electron density, or into plausible interfacial positions in both the upper and lower leaflets, failed to prevent deformation of the TMD (Supplementary Figure S11). With or without manually placed lipids, lipids frequently penetrated the TMD-subunit interface, in some cases entering the channel pore (Supplementary Figure S12). The associated deformations were further associated with variable pore collapse (Figure 4G; Supplementary Figure S7A), rendering this system unsuitable for open-state simulations. Thus, whereas closed-pore structures were largely validated by MD, at least in our hands the wide-open X-ray state did not appear to be stable under any simulation conditions attempted.

**Figure 4:**
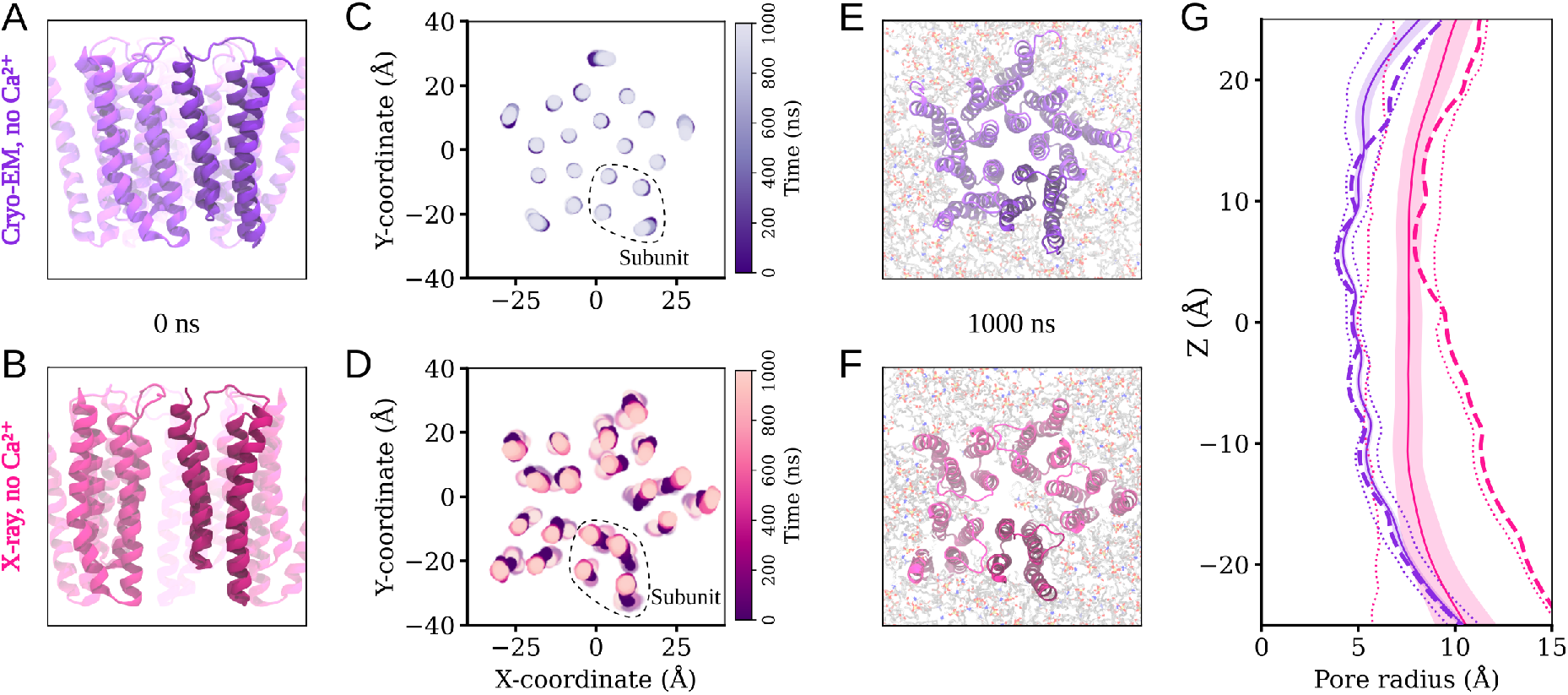
Behaviour of the transmembrane domain in molecular dynamics simulations. (A,B) View of the transmembrane region of DeCLIC, seen from the membrane plane, prior to MD simulation in the calcium-free cryo-EM structure and in the calcium-free X-ray structure. (C,D) Position of the center of mass of each transmembrane helix in the bilayer plane through 1000 ns of MD simulations and four replicas. For the calcium-free cryo-EM structure the helices of the transmembrane region remain close to their positions in the structure, while the transmembrane domain in simulation of the X-ray structure without calcium undergo a conformational change to an asymmetric arrangement of the subunits. (E,F) The transmembrane region seen from the extracellular side at the end of the simulations. The cryo-EM structure without calcium has retained a symmetric arrangement, while the X-ray structure without calcium has become asymmetric, with lipids penetrating to the pore in one simulation replica. (G) Pore profiles tracing the protein backbone for the calcium-free cryo-EM (purple) and X-ray (pink) systems. Dashed lines show the profile for the starting structure, solid lines the simulation average with standard deviation in colored fill, and minimum and maximal values during the simulations in dotted lines. There is little variation in the pore profile of the cryo-EM system during the simulations, while the wide lower part of the pore in the calcium-free X-ray structure contracts during the simulations.

### Simulations of alternative NTD states refine fit to SANS data

In contrast to the ECD and TMD, in all simulation systems the NTD was notably dynamic, moving substantially relative to its starting conformation (Figure 5A–B; Supplementary Figure S13-S15). Indeed, when aligned on the ECD-TMD scaffold, average C_*α*_ RMSDs for the NTD1 and NTD2 lobes reached over 10 Å and 6 Å respectively (Supplementary Figure S9B–C). This variability could largely be attributed to rigid-body motions of individual lobes, as deviations within each lobe remained at 3 Å when it was aligned on itself (Supplementary Figure S9B–C). The NTD1 lobes sampled a range of positions, often translated outward from the pore (z-)axis and down towards the membrane; NTD2 primarily translated up or down along z (Figure 5A–B; Supplementary Figure S13-S15). In several cases, these transitions appeared to be reversible, with lobes returning to positions near their starting poses within 1 µs (Supplementary Figure S14–S15).

**Figure 5:**
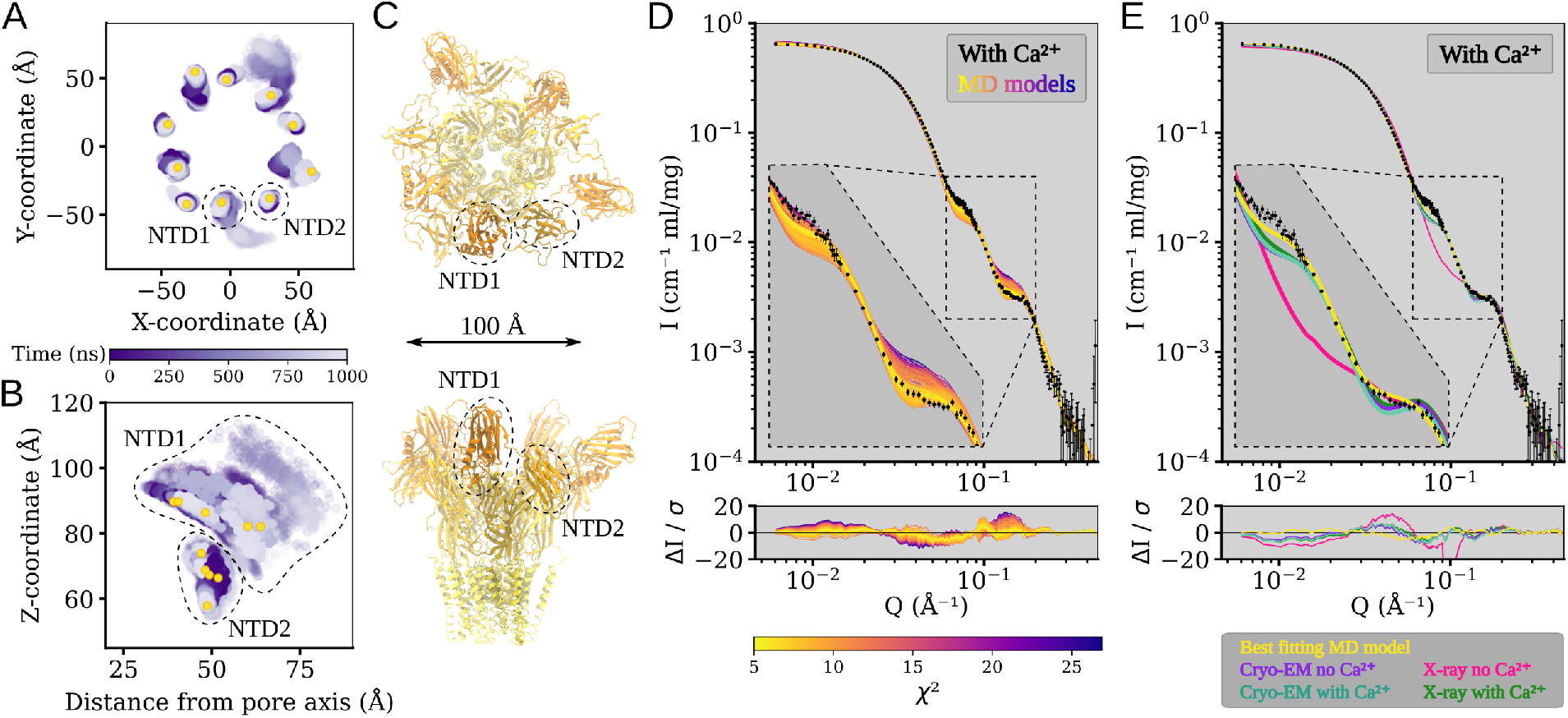
The N-terminal domains of DeCLIC are highly dynamic, and an asymmetric arrangement thereof with a mix of contracted and extended NTD positions yield the best fit to the SANS data. (A,B) Position of the center of mass of each NTD lobe over time in four MD simulations, showing their positions in the XY-plane (A) and their Z-coordinate as a function of distance from the pore axis (B). NTD1 is highly mobile, sampling positions further out, lower down, and higher up than the starting conformation. NTD2 has less mobility, mainly sampling positions along Z. Yellow dots show the position of the NTD lobes in C. (C) Snapshot of the MD simulation frame yielding the best fit to the with-calcium SANS data (χ^2^ of 5.2), seen from the extracellular side (top) and from the plane of the membrane (bottom). The protein has adopted an asymmetric conformation with three NTD1 domains in positions similar to the determined structures and two NTD1 further out and down. The snapshot is from the simulations started from the no-Ca^2+^ cryo-EM structure. (D) Fits of snapshots from simulations of the no-Ca^2+^ cryo-EM structure and the error weighted residual between the models and the scattering profile. The simulations yield theoretical distributions around the experimental scattering curve, containing models with better fit (best χ^2^ of 5.2) to the data than the experimental structure from which the simulations were launched (χ^2^ of 11.2). Inset shows *Q*∈[0.06, 0.2] Å ^-1^. (E) Comparison between the best fitting model from simulations (yellow), structures of closed-like DeCLIC (purple, aquamarine, and green; note that curves overlap), and the X-ray no-Ca^2+^ structure (pink). As seen in the inset showing *Q ∈* [0.06, 0.2] Å ^-1^, the simulation model has the best fit to both features in the scattering profile.

All simulations included snapshots that improved goodness of fit to our Ca^2+^-SANS data, relative to our static cryo-EM or X-ray structures (Figure 5D, Supplementary Figures S16-S17). Better goodness of fit to the feature at *Q* [0.06, 0.09] Å ^-1^ generally corresponded to alternative conformations of the NTD, particularly asymmetric arrangements of NTD1. For instance, in the overall best-matching snapshot (χ^2^ = 5.2; Figure 5C, E) two NTD1 lobes were translated outward from the z-axis (i.e. the pore axis) and down towards the membrane, relative to the starting model (cryo-EM model without calcium, χ^2^ = 11.2). Other snapshots produced even better goodness of fit to the *Q*∈ [0.06, 0.09] Å ^-1^ feature, with more dramatic translation of the NTD1 lobes (Supplementary Figure S18). However, SANS curves calculated from these models were poor matches for the *Q*∈[0.11, 0.19] Å ^-1^ feature that corresponds to shorter interatomic distances, resulting in worse goodness of fit overall. Interestingly, simulations did not improve goodness of fit to our *apo*-SANS data: the best simulation frame (χ^2^ = 9.4) was comparable to the best static model (X-ray model with calcium, χ^2^ = 8.8) (Supplementary Figures S16-S17), and while it did improve correspondence to long pairwise distances, it did not match the distribution of medium distances (Supplementary Figure S2B). Thus in the presence of calcium, DeCLIC’s solution state was best approximated by an asymmetric modification of the cryo-EM structure, including some NTD1 lobes translated “out-and-down”. In the absence of calcium, no single structure or snapshot provided as strong a prediction of the SANS profile, possibly implicating the presence of an alternative state.

## Discussion

The combination of SANS, cryo-EM, and MD has enabled us to describe a plausible mechanism for extracellular calcium modulation and ion permeation, and of dynamic NTD states corresponding to solution-phase conditions (Figure 6). When calcium is present, it is tightly bound at an interfacial site in the canonical ECD, where it blocks ions from transiting between the extracellular vestibule and external medium via this interface (Figure 6, *center*). In the absence of calcium, monovalent cations such as sodium can traverse this interfacial site, possibly facilitated by the electronegative cluster of glutamate residues on the principal *β*1–*β*2 loop and complementary loop F (Figure 6, *right*). This phenomenon may be particularly relevant to ion permeation, given the presence of a constriction in the outer ECD vestibule (Supplementary Figure S7), which could limit transit to or from the transmembrane channel via a linear pathway. Notably, ion paths to the transmembrane pore through electronegative fenestrations rather than the linear axis of the ECD have also been reported in both nicotinic [14] and GABA_A_ receptors [15–17], including acidic residues on the *β*1–*β*2 loop, Cys-loop, and loop F.

**Figure 6:**
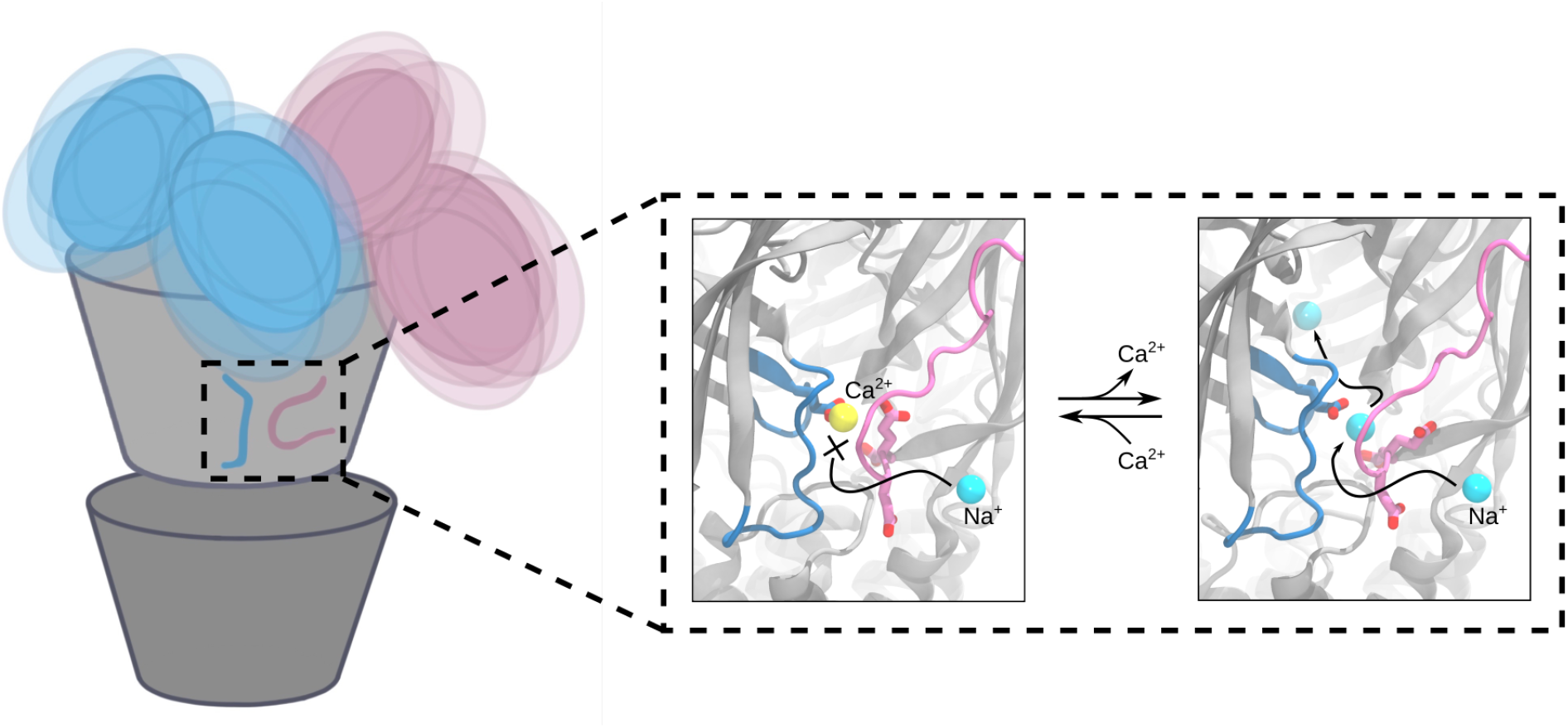
Conformational variability and proposed calcium binding site behaviour in closed DeCLIC. The positions of the N-terminal domains of DeCLIC fluctuate around the core of the protein, sampling a wide range of conformations where the average can be described by an asymmetric conformation with a mix of compact and extended NTD positions. In the calcium binding site, bound calcium blocks sodium access to the site. Following calcium depletion, sodium can enter the central pore through the calcium binding site. The calcium binding site is thus proposed to act as a supplementary way for sodium ions to reach the ion conduction pathway, with the three consecutive glutamates on loop F shepherding sodium from outside the protein, through the calcium binding site, and into the extracellular part of the ion channel pore.

These effects are further reminiscent of calcium modulation in the larger family of pLGICs, where calcium has been shown to inhibit both the bacterial ELIC and eukaryotic serotonin receptor (5-HT_3A_R) [10, 18], or to potentiate neuronal *α*7-nicotinic acetylcholine receptors (nAChR) [19, 20]. Divalent zinc ions have also been shown to inhibit GABA_A_ receptors [21], and to exert bimodal effects on glycine (GlyR), nicotinic, and 5-HT_3A_ receptors [22–24]. Although a variety of specific residues have been implicated in these effects, most are located at the ECD subunit interface, involving loop F as well as the *β*1–*β*2 and Cys/Pro-loops (Figure 7) [10]. Acidic loop-F residues E150 and D158 in ELIC [10], E213 and E215 in 5-HT_3A_R [25], E172 in *α*7-nicotinic [26, 27], and E182 in GABA_A_ receptors [28] have been shown to be particularly important, suggesting a common strategy for divalent ion modulation.

**Figure 7:**
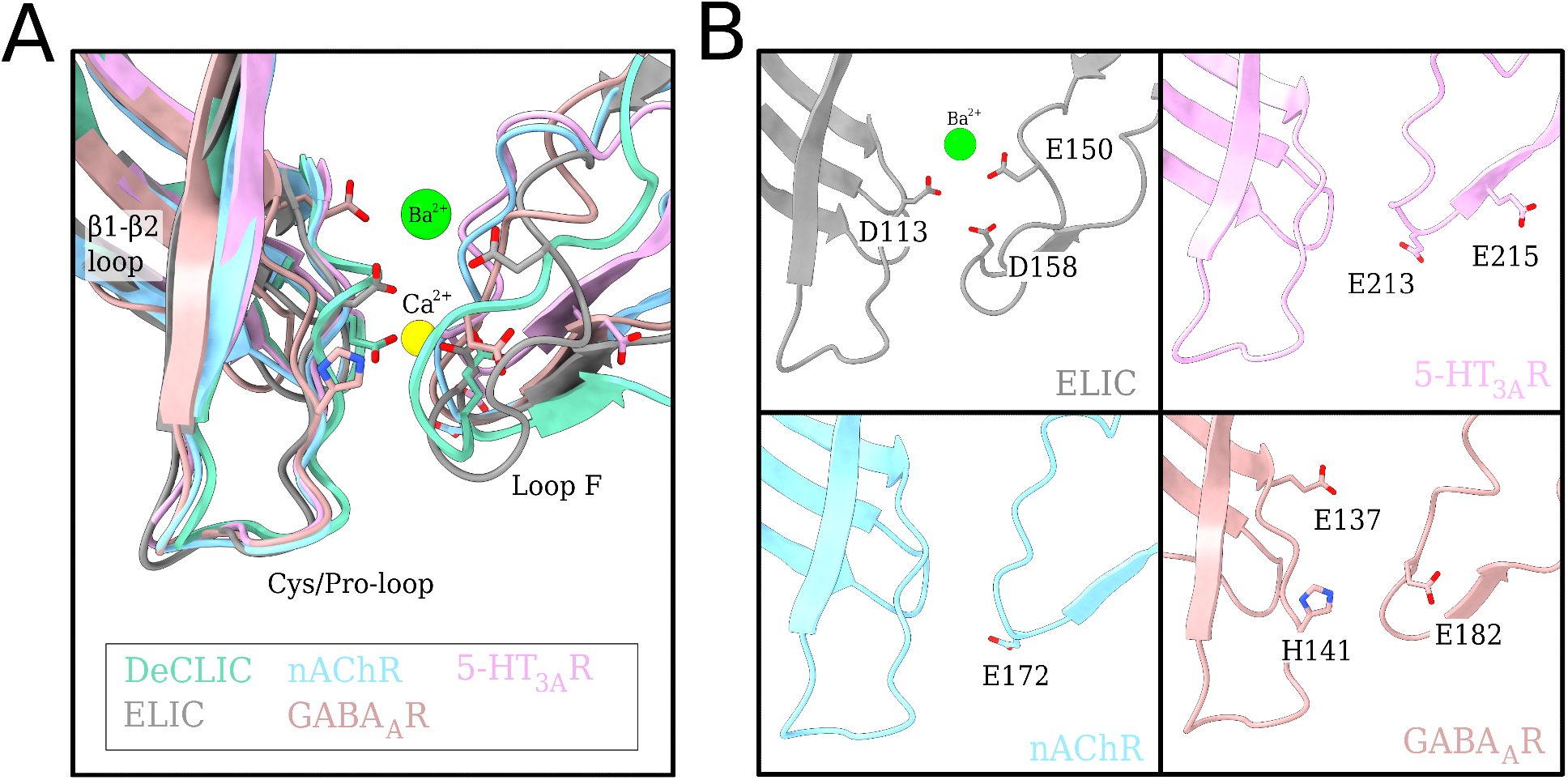
Charged residues at the subunit interface present in other family members. (A) Overlay of the subunit interface (loop F, Cys/Pro-loop and *β*1-*β*2 loop) from DeCLIC with other family members nAChR (dark blue), ELIC (grey), 5-HT_3A_R (dark grey) and GABA_A_R (purple) with Ca^2+^ ion in yellow and Ba^2+^ in green. The important charger residues are displayed as sticks (colored by heteroatom). (B) Display of each family member from (A) individually for clearer representation with relevant charged residues displayed as sticks and colored by heteroatom.

Aside from the ECD calcium site, structures in the presence and absence of calcium were largely superimposable, including a closed pore. This observation was initially surprising, given previous evidence for channel opening upon calcium depletion, from both electrophysiology and crystallography [11]. It is possible that only a minority population of channels is open even under maximally conducting conditions. Alternatively, DeCLIC may require an additional signal aside from calcium depletion to open substantially under SANS or cryo-EM conditions. It is also theoretically possible that a fraction of open channels exist in the data, but that particle classification failed to identify them as a substantial discrete population of channels, as has previously been reported for the related bacterial channel GLIC [29].

Pairwise distance distributions from SANS data did indicate the promotion of an alternative state in the absence of calcium, consistent with a limited population similar to the open X-ray structure. Similarly, the radius of gyration was moderately reduced by calcium removal (Table S1, *multiple methods*), indicating a contribution of relatively compact NTD states under *apo* conditions. Further rigidification of the NTD—for example, via crystal contacts—could stabilize an open state of the channel, otherwise underrepresented in solution. However, MD simulations were capable of improving goodness of fit to the SANS profile in calcium but not *apo* conditions, indicating that the available structures—closed and open—did not spontaneously sample conformations representative of the *apo* time and population average. We have previously reported improved SANS goodness of fits to specific pLGIC states by linear combination of closed and open structures [6]; however, such an approach relies on trusted models of all conformations contributing to the population. Instead, here our simulations failed to support the open X-ray structure as a stable state, leaving open questions as to the physiologically relevant conductive state.

Although both X-ray and cryo-EM structures substantiated a symmetrical, compact predominant state of the DeCLIC NTD, data from all three methods here also supported substantial mobility in this domain (Figure 6, *left*). Flexibility in the NTD is consistent with its relatively low resolution, and with motions directly observed in MD simulations, as well as the ability of MD results to describe solution-phase SANS data better than static structures. Because SANS profiles reflect the time- and population-average conformation, this model could either represent the most probable conformation of an individual protein, or an average of a mixture of states. Simulation results offer more specific predictions that asymmetric arrangements of the NTD lobes are common and rapidly interchangeable. The range of NTD motion allowed this domain to sample a volume around the protein considerably larger than originally predicted by structural data, possibly facilitating encounters between the NTD and yet-unknown interaction partners in the periplasmic space. Thus, alongside insights into ion interactions and pore stability, this work highlights conformational features and intrinsic challenges of determining structures of proteins with differentially dynamic domains.

## Materials and Methods

### DeCLIC Expression and Purification

Expression and purification of DeCLIC-MBP was adapted from the protocol published by Hu et al. [11]. In short, C43(DE3) *E. coli* transformed with DeCLIC-MBP in pET-20b were cultured overnight at 37° C. Cells were inoculated 1:100 into 2xYT media with 100 µg/mL ampicillin, grown at 37° C to OD_600_ = 0.8, induced with 100 µM isopropyl-β-D-1-thiogalactopyranoside (IPTG), and shaken overnight at room temperature. Cells were resuspended in buffer A (300 mM NaCl, 20 mM Tris-HCl pH 7.4) supplemented with 1 mg/mL lysozyme, 20 µg/mL DNase I, 5 mM MgCl_2_, and protease inhibitors. Membranes were harvested from cell pellets by sonication and ultracentrifugation, then immediately solubilized in 2% n-dodecyl-β-D-maltoside (DDM). Fusion proteins were purified in batch by amylose affinity (NEB), eluting in buffer B (buffer A with 0.02% DDM) with 2–20 mM maltose, then further purified by size exclusion chromatography (SEC) in buffer B. After overnight thrombin digestion, DeCLIC was isolated from its fusion partner with a final size exclusion run, and concentrated to 3–5 mg/mL by centrifugation.

### Small-Angle Neutron Scattering

SEC-SANS experiments [30] were performed at the D22 beamline of Institute Laue–Langevin, using a paused flow approach [6]. In short, the protein sample was loaded on a Superdex 200 Increase 10/300 gel filtration column in-line with the SANS measurement[31, 32], exchanging the sample to the D_2_O buffer environment and match-out deuterated DDM (d-DDM) [33] prior to the protein reaching the SANS measuring cell. Upon peak detection by UV-vis absorvance at 280 nm, the flow was slowed to 0.01 ml/min, and at the peak max the flow was paused. Two detector distances were used to measure the scattered intensity as a function of momentum transfer Q, 2.8m/2.8m during the run, and 8m/8m while paused. The definition of Q used was Q=(4π/λ)sin(*θ*), where 2*θ* is the scattering angle and λ is the wavelength. Measurements were performed under two buffer conditions, with calcium (D_2_O, 150 mM NaCl, 20 mM Tris*·*HCl, 10 mM CaCl_2_, 0.5 mM d-DDM) and without calcium (D_2_O, 150 mM NaCl, 20 mM Tris HCl, 10 mM EDTA, 0.5 mM d-DDM).Further sample details and collection parameters are available in Supplementary Tables S5 and S6.

Data reduction was performed using Grasp version 9.04 [34], correcting for the empty cuvette and background, scaling by transmission and thickness, scaling to absolute intensity by direct flux measurement, and subtracting the buffer contribution using data from a dedicated buffer measurement. Concentration normalization was performed by dividing the scattering intensity with the average protein concentration during the measurement, which was calculated from the co-recorded chromatogram and the extinction coefficient calculated from the amino acid sequence using ProtParam [35]. Data from the two detector distances were merged, using data collected at 8m up to 0.09 Å ^-1^, and for higher Q-values data collected at 2.8m. A small additional constant was subtracted as a final adjustment of the background.

Guinier analysis was used to calculate I(0) and radius of gyration (R_g_), and the molecular weight was estimated from I(0), see Supplementary Table S1. The excess scattering length density was calculated using the scattering length density of DeCLIC and D_2_O, see Supplementary Tables S5, accounting for hydrogen-deuterium exchange of labile hydrogens in DeCLIC by calculating the expected degree of exchange after 100 minutes (approximately the time from the start of a run until the peak max) at pH 7.5 using PSX (Protein-Solvent Exchange) [36]. As DeCLIC is a transmembrane protein the labile hydrogens in the transmembrane region can be expected to be shielded from exchange, which was accounted for by considering hydrogens withing the transmembrane region, as defined by the Positioning of Proteins in Membranes (PPM) webserver [37], as ^1^H. The partial specific volume was estimated using the volume calculated by ^3^V [38] for the X-ray structure with calcium, and the molecular weight expected from the amino acid sequence. Pair distance distributions from the experimental SANS were calculated using BayesApp [39] – available at https://somo.chem.utk.edu/bayesapp/ – and compared with distribution calculated with CaPP (Calculating Pair distance distribution functions for Proteins) [40] from all atom models. Theoretical scattering curves from all atom models (cryo-EM structures, X-ray structures, molecular dynamics snapshots) and fits of them to the experimental scattering profiles were calculated using Pepsi-SANS [41]. A summary of equations and software used is available in Supplementary Table S7.

### Cryo-EM Sample Preparation and Data Acquisition

For each of the experimental conditions, 3 µl of DeCLIC sample was applied to a glow-discharged quantifoil 1.2/1.3 Cu 300 mesh grid (Quantifoil Micro Tools), which was then blotted for 1.5 s and plunge-frozen into liquid ethane using a FEI Vitrobot Mark IV. Micrographs were collected on an FEI Titan Krios 300 kV microscope with a post energy filter Gatan K2-Summit direct detector camera. Movies were collected at nominal 165,000x magnification, equivalent to a pixel spacing of 0.82Å. A total dose of 40 e^-^/Å ^2^ was used to collect 40 frames over 6 sec, using a nominal defocus range covering -1.4 to -3.2 µm (for the no-Ca^2+^ dataset) and -2.0 to -3.6 µm (for the Ca^2+^-present dataset) in steps of 0.2 µm. There is a slight difference in the defocus values of the collected datasets, which was selected to optimize the signal-to-noise ratio and contrast of the particles for collection on two different microscopes.

### Image Processing

Motion correction was carried out with MotionCor2 [42]. All subsequent processing was performed through the Relion 3.1 pipeline [43]. Defocus was estimated from the motion corrected micrographs using CtfFind4 [44]. Following manual picking, initial 2D classification was performed to generate references for autopicking. Particles were extracted after autopicking and binned before the initial reference was generated. The aberrant particles were removed from the dataset through multiple rounds of 2D- and 3D-classification as well as 3D auto-refinement. Per-particle CTF parameters were estimated from the resulting reconstruction using Relion 3.1. Global beam-tilt was estimated from the micrographs and correction applied. The final 3D auto-refinement was performed using a soft mask, followed by post-processing with the same mask. Local resolution was estimated using the Relion implementation. Post-processed densities were improved using ResolveCryoEM, a part of the Phenix package (release 1.18 and later) [45] based on maximum-likelihood density modification [46]. Densities from both Relion post-processing and ResolveCryoEM were used for building; figures show output from Relion post-processed (Fig 3, Fig S6).

### Model Building

Models were built starting from a monomer of an X-ray structure determined at pH 7 in the presence of Ca^2+^ (PDB ID: 6V4S [11], chain A), where the missing residues in the NTD region were built by Modeller [47]. Phenix 1.19.2-4158 [45] real-space refinement was used to refine the model, imposing 5-fold symmetry through NCS restraints detected from the reconstructed cryo-EM map (Correlation of symmetry-related regions: 0.96 and 0.97 for the two reconstructions). The model was manually adjusted in Coot 0.8.9.1 EL [48] and re-refined until the quality metrics were optimal and in agreement with the reconstruction. Model statistics are summarized in Table S3. Additionally, the side chains and residues, where the density did not allow for confident building, were removed (Table S4).

### Molecular Dynamics Simulations

Molecular dynamics simulations were performed starting from the cryo-EM and X-ray structures resolved in the presence (PDB ID:s 7Q3G and 6V4S) and absence of calcium (PDB ID:s 7Q3H and 6V4A). For the X-ray structures missing residues were built using Modeller v. 9.22 [47], for the cryo-EM structures the full-length models prior to removal of residues with insufficient density were used for simulation. Protonation states were set to neutral pH using Pdb2Pqr [49] or with Charmm-GUI [50, 51], with equivalent result. The ions resolved in the with-calcium structures were retained for the simulations, additionally a system was prepared from the with-calcium X-ray structure where the resolved ions were removed. For the X-ray structure resolved in the absence of calcium, three simulation systems were prepared; one without lipids intercalated between the subunits, one where an upper leaflet 1-palmitoyl-2-oleoyl-sn-glycero-3-phosphocholine (POPC) lipid was manually placed guided by the intercalated head-group resolved in structure, and one with both an upper and a lower leaflet POPC lipid manually intercalated between subunits. Simulation systems were set-up using the Charmm-GUI membrame builder [50, 51]; embedding the protein in a POPC bilayer, adding sodium and chloride ions to neutralize the system and to 150 mM NaCl concentration, and solvating in TIP3 water. Hydrogen mass re-partitioning was used for the simulations starting from the cryo-EM structures. The CHARMM36m force-field was used [52], and simulations performed using Gromacs-2020.3 [53]. Systems were energy minimized, followed by equilibration for 2.25 ns during which restraints were gradually released. Four replicate production simulations of 1000 ns were performed for each system.

Root mean square deviation (RMSD) and root mean square fluctuations (RMSF) were calculated using Gromacs. RMSDs were calculated for C-alpha atoms of full subunits and for individual structural domains, using the selection itself to align the trajectories and aligning on the ECD together with the TMD. RMSF was calculated for loop F, including the full residues and aligning on loop F. Distances and center of mass positions were tracked through the trajectories using Vmd [54]. Pore profiles for the simulations and for the structures were calculated using Chap [55]. Vmd [54] and UCSF Chimera [56] were used to render images.

## Data Availability

Cryo-EM density maps of the pentameric ligand-gated ion channel DeCLIC in detergent micelles have been deposited in the Electron Microscopy Data Bank under accession numbers EMD-13791 (pH 7, 10 mM Ca^2+^) and EMD-13792 (pH 7, no Ca^2+^). Each deposition includes the cryo-EM sharpened and unsharpened maps, both half-maps and the mask used for final FSC calculation. Coordinates of the two models have been deposited in the Protein Data Bank. The accession numbers for the two DeCLIC structures are 7Q3G (pH 7, 10 mM Ca^2+^) and 7Q3H (pH 7, no Ca^2+^). The raw experimental SANS data is available from https://doi.ill.fr/10.5291/ILL-DATA.8-03-1002, and processed SANS data – with best fitting models for the with-calcium data-set – are available in the Small Angle Scattering Biological Data Bank (SASBDB) as entries SASDNG5 and SASDNH5. MD simulation trajectories are available at Zenodo.org as entry https://doi.org/10.5281/zenodo.6022369.

## Acknowledgments

The authors would like to thank the Swedish Cryo-EM National Facility staff, especially Marta Carroni and Stefan Fleischmann from Stockholm and Michael Hall from Umeå, for kind assistance with data collection. We would also like to acknowledge Institut Laue-Langevin for development of SEC-SANS and allocated beam time at the D22 instrument, and in particular Lionel Porcar for assistance with data collection. This work was supported by the Swedish Foundation for Strategic Research through the SwedNess graduate school (GSn15- 0008), the Knut and Alice Wallenberg Foundation, and the Swedish Research Council (2019-02433, 2021- 05806), the Erasmus plus program (AH), the Swedish e-Science Research Centre, and the BioExcel Center of Excellence (EU-823830). Cryo-EM data were collected at the Swedish national cryo-EM facility funded by the Knut and Alice Wallenberg Foundation, Erling Persson and Kempe Foundations. Computational resources were provided by the Swedish National Infrastructure for Computing (SNIC).

## Author Contributions

Conceptualisation: RJH, EL; methodology: ML, UR, RJH; software: ML, UR, AH; validation: ML, UR, AM; formal analysis: ML, UR, AH, AM; investigation: ML, UR, RJH; resources: AM, RJH, EL; data curation: ML, UR, RJH, EL; original draft: ML, UR, RJH; review and editing: ML, UR, AH, AM, RJH, EL; visualization: ML, UR; supervision: RJH, EL; project administration: AM, RJH; funding acquisition: EL.

### Competing interests

The authors declare that they have no conflict of interest.

## Supplementary Information

**Figure S1:**
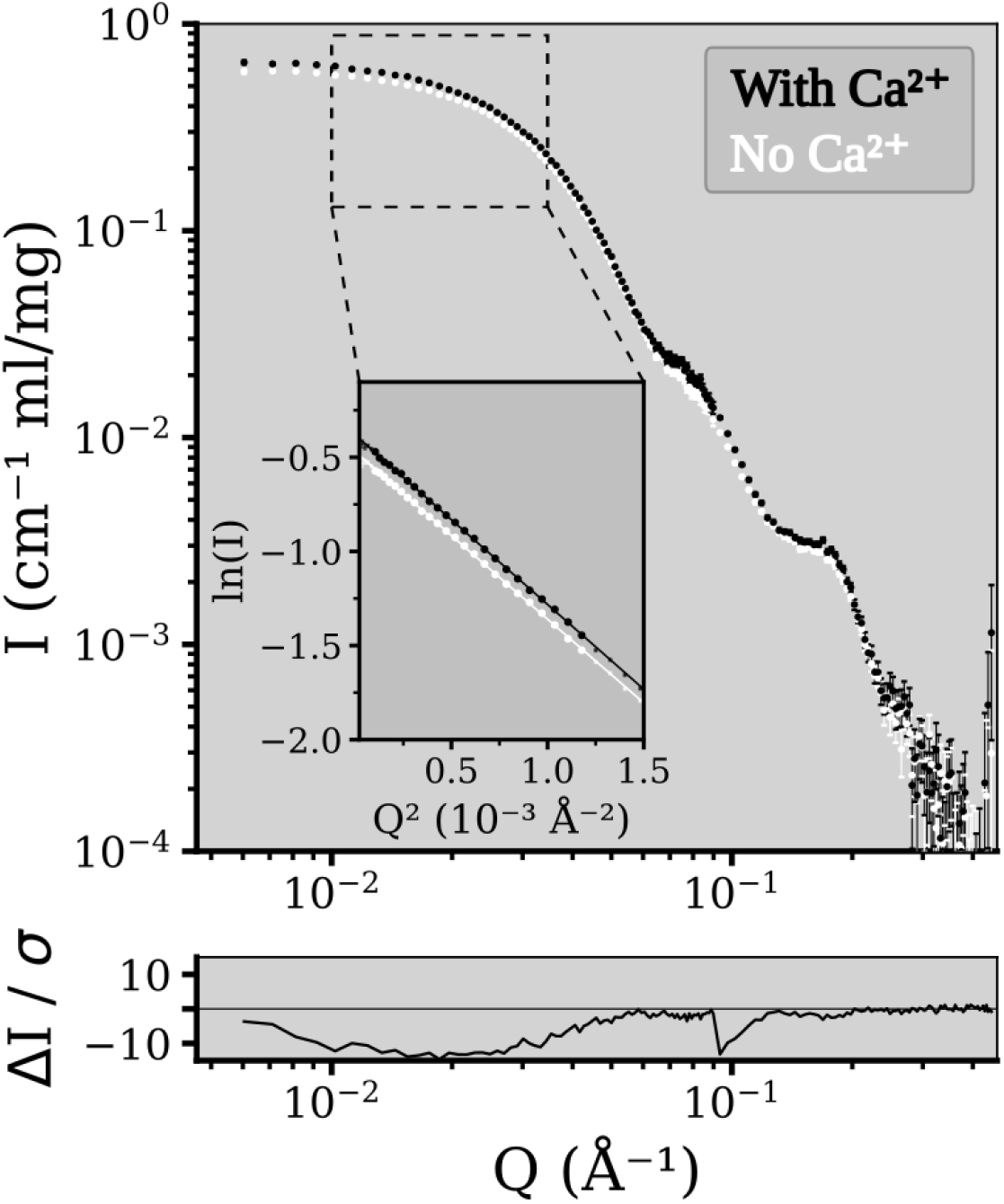
Comparison of small angle neutron scattering from DeCLIC in the presence and absence of calcium. Small angle neutron scattering profiles from DeCLIC in the presence (black) and absence (white) of calcium, with the insert showing the Guinier plot of the low Q region together with the Guinier fits as lines. The lower panel shows the error weighted residual between the conditions.

**Figure S2:**
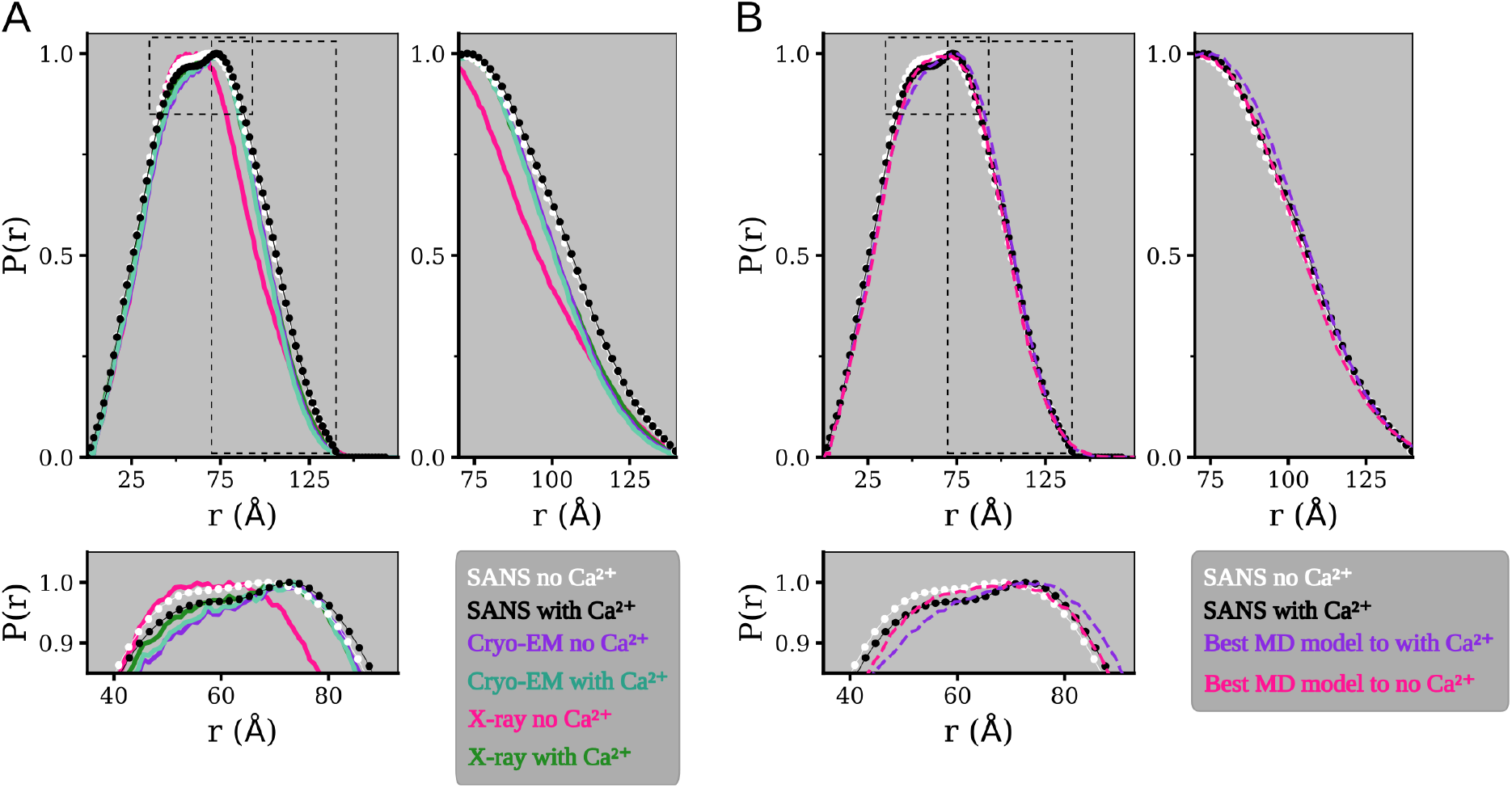
Pairwise distance distributions from SANS compared to that of structures and models. (A) Pairwise distance distribution from SANS data with (black) and without (white) calcium, and for structures from cryo-EM and X-ray crystallography determined in the presence (aquamarine for cryo-EM, green for X-ray) and absence of calcium (purple for cryo-EM, pink for X-ray). The lower box shows a zoom on the peak of the distributions, the box to the right shows a zoom on the larger distances. The peak show that SANS in the absence of calcium yields a distribution with features similar to both the X-ray no Ca^2+^ structure and to the closed-like structures, while the SANS data with Ca^2+^ is closest to the X-ray with calcium structure. The long distances show that the structures in this region underestimate the values observed with SANS. (B) Pairwise distance distribution from SANS data with (black) and without (white) calcium, and for the models from molecular which gives the best fit to the SANS data. The purple model gives the best fit to the with Ca^2+^ SANS data and comes from the simulations of the calcium free cryo-EM structure. The pink model gives the best fit to the no Ca^2+^ SANS data and comes from the simulations of the calcium free X-ray structure. The lower box shows a zoom on the peak of the distributions, the box to the right shows a zoom on the larger distances. It can be seen in the peak region that these models are not perfect matches for the distance distributions obtained from SANS, while the larger distances show that the agreement between the models and the SANS data has improved in this region.

**Figure S3:**
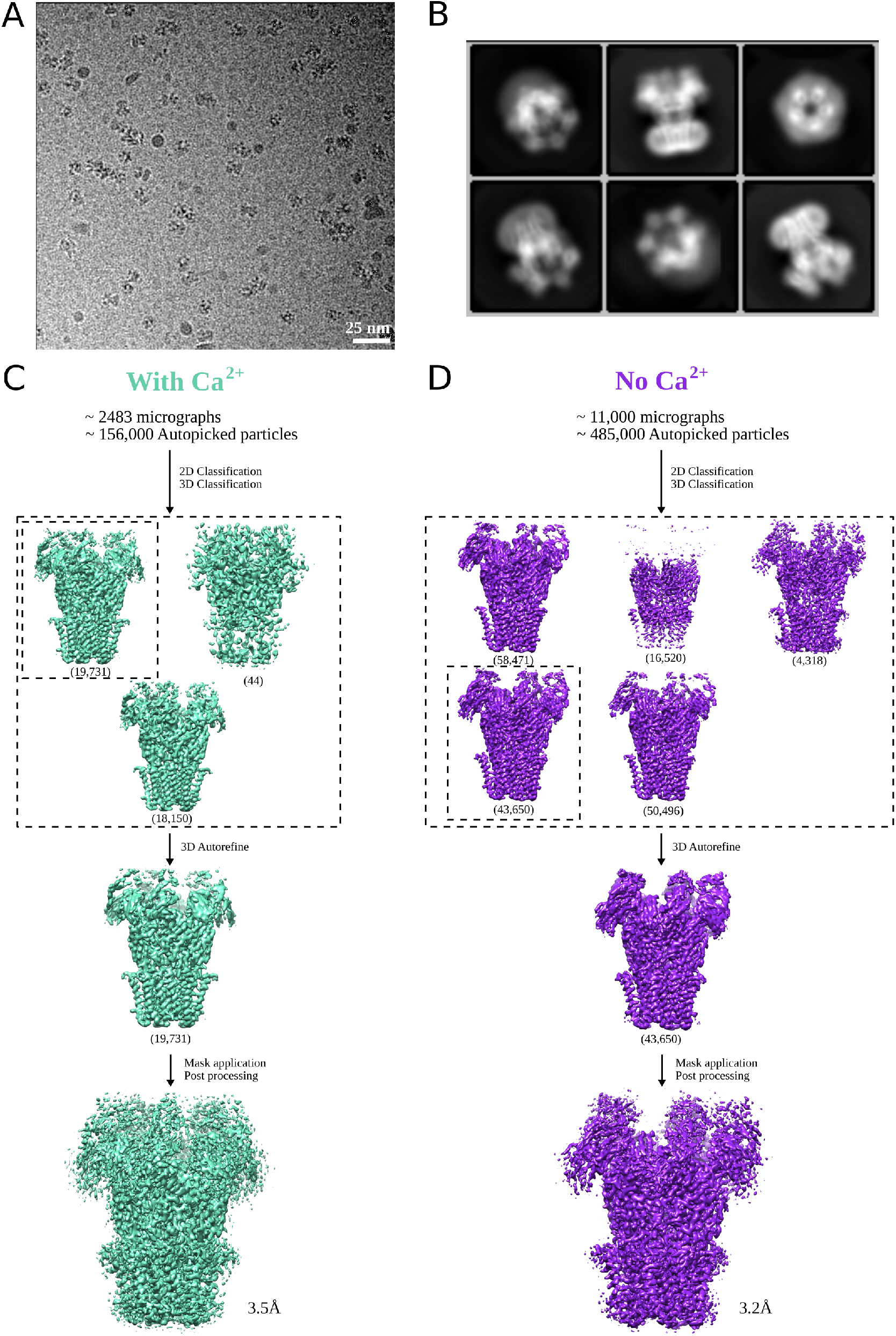
Cryo-EM data and processing. (A) Representative micrograph from one of the grids used for data collection on a Titan Krios, showing detergent-solubilized DeCLIC particles. (B) Representative 2D class averages at 0.82 Å /px in a 256 × 256 pixel box and a 200 Å mask. (C) Overview of the cryo-EM processing pipeline for data collected at pH 7 with Ca^2+^. (D) Same as in C but for a dataset where no Ca^2+^ was present (see Methods).

**Figure S4:**
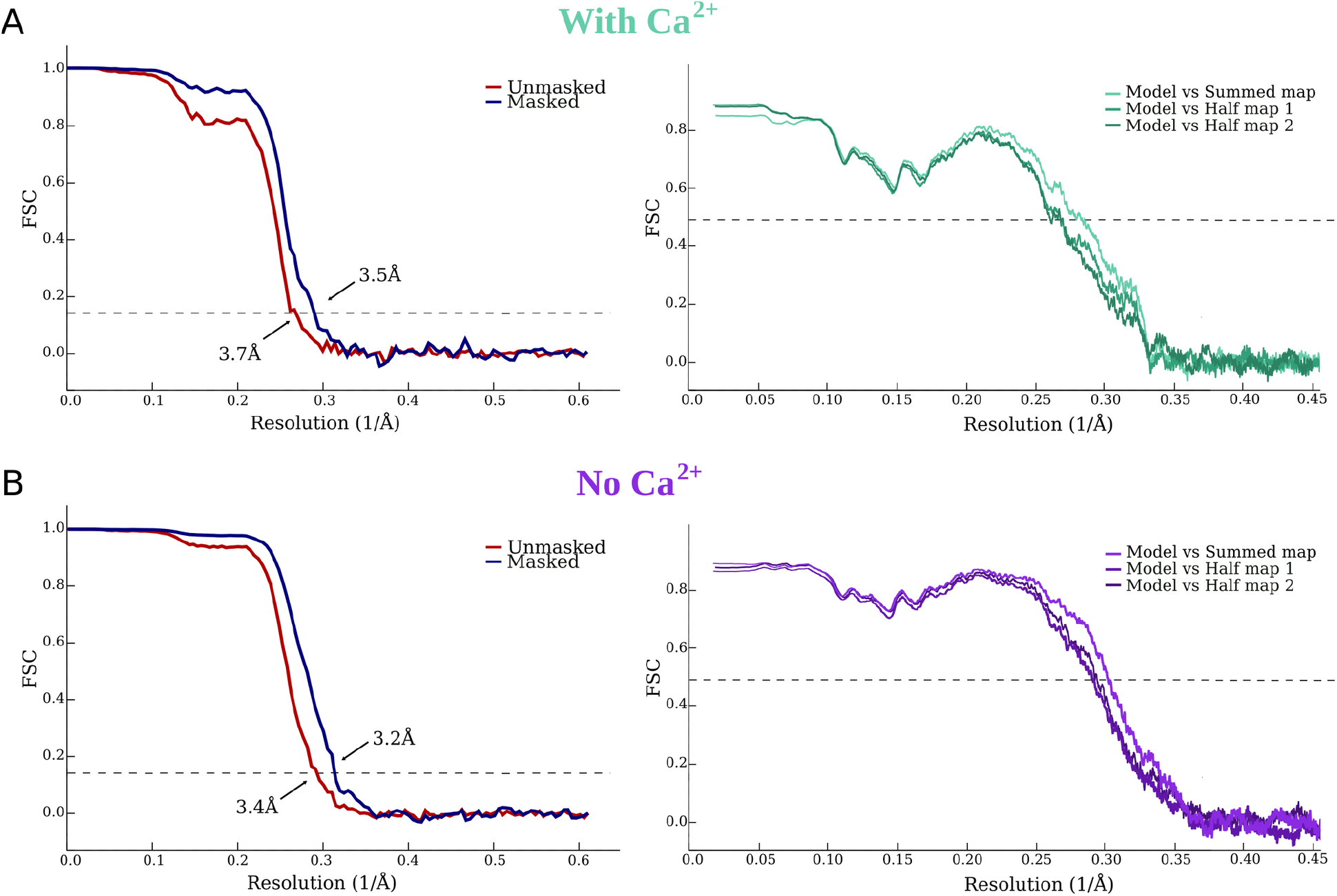
Fourier Shell Correlation. (A) *Left* : FSC curves for cryo-EM reconstructions at pH 7 with Ca^2+^ before (red) and after (blue) applying a soft mask. *Right* : FSC curves for cross-validation between model and half-map 1 (medium), model and half-map 2 (dark) and model and summed map (light). (B) FSC curves represented as in panel A for the dataset at pH 7 without Ca^2+^.

**Figure S5:**
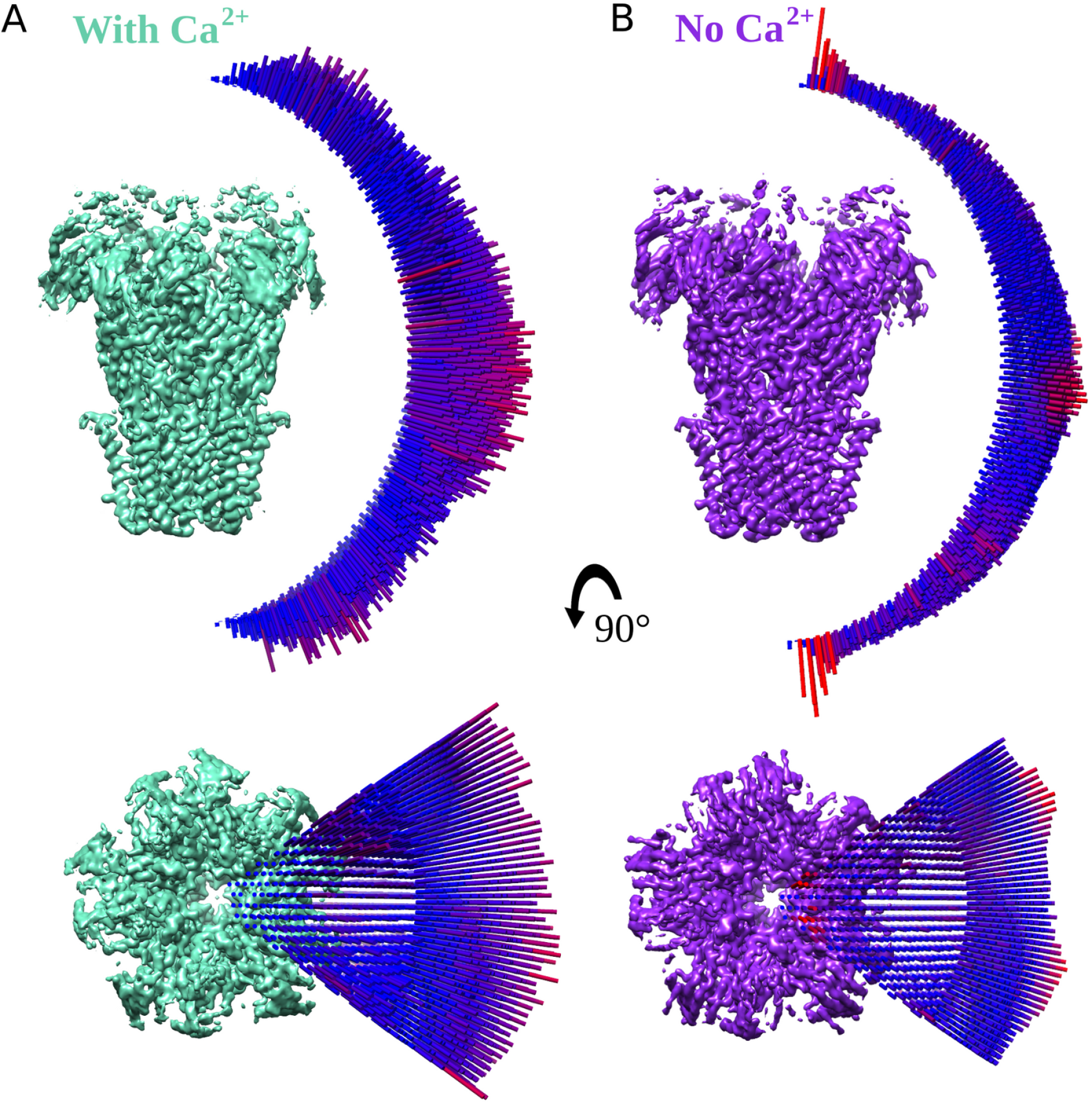
Distribution of orientation sampling in the cryo-EM maps. (A) *Top*: Side view of the cryo-EM map for DeCLIC at pH 7 with Ca^2+^ and the distribution of orientation sampling after refinement. The height and the color (blue to red) of the bar represent the abundance of that particular view. *Bottom*: Representation of the same map from the top. (B) Map and distributions as in panel A for the dataset at pH 7 with no Ca^2+^.

**Figure S6:**
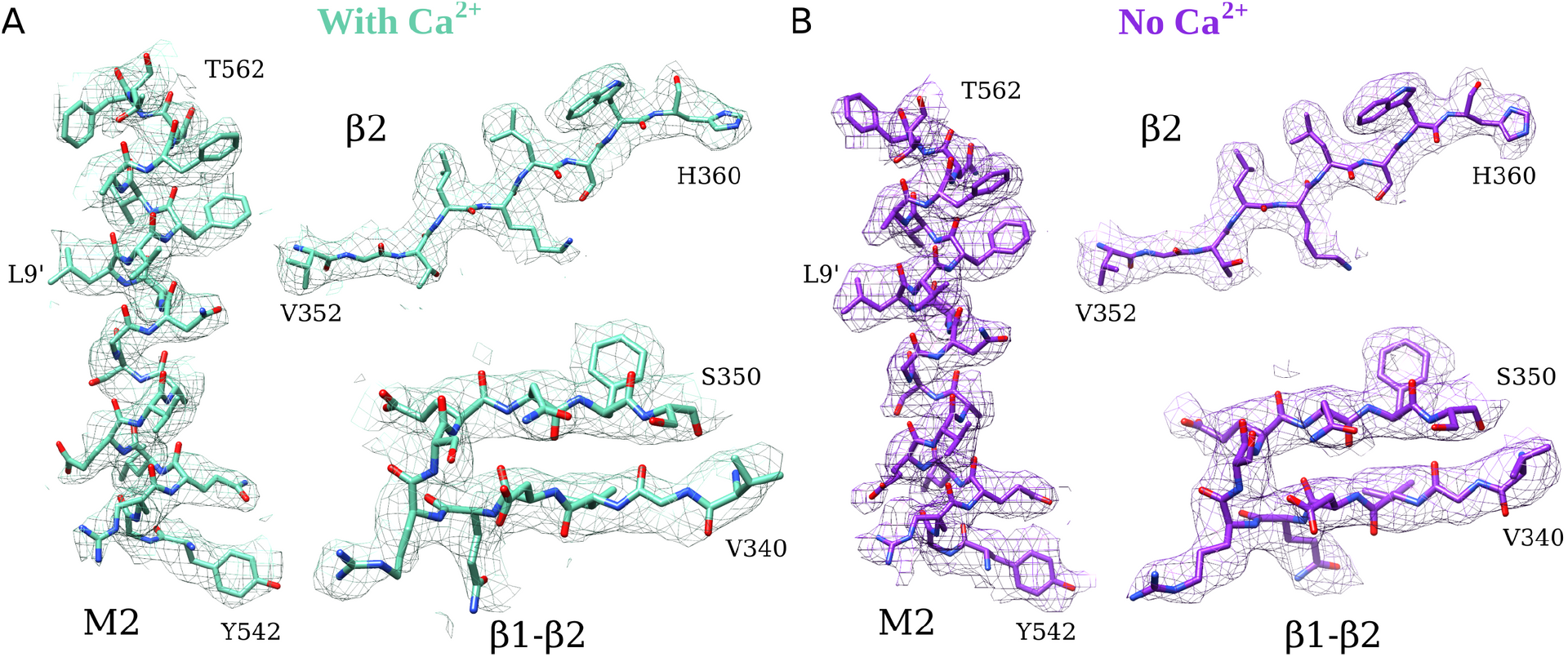
Cryo-EM map and atomic model for select regions of DeCLIC. (A) Density (mesh) and corresponding atomic model (sticks, colored by heteroatom) for the M2 helix (Y542–T562), part of the β2 strand (V352-H360) and a β1-β2 loop (V340-S350) for the pH 7 with Ca^2+^ model. (B) Density and corresponding model, shown as in panel A, for the pH 7 model without Ca^2+^.

**Figure S7:**
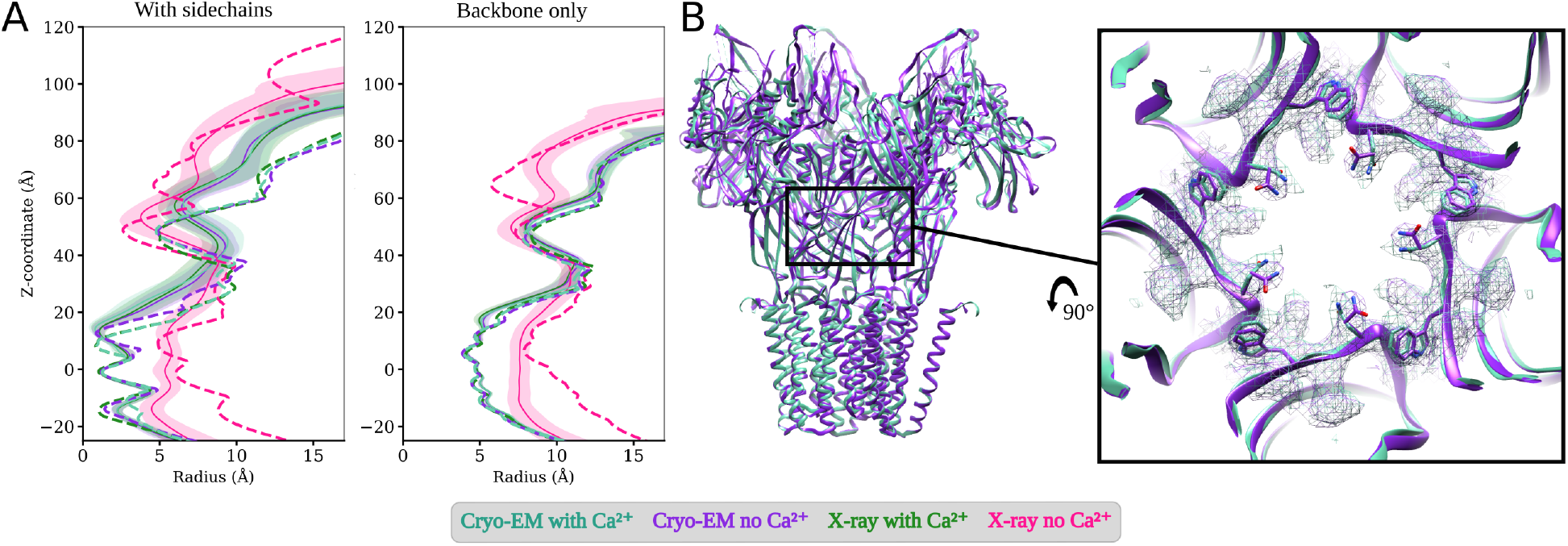
Pore profiles of DeCLIC and zoom in view of the constriction point in the ECD. Pore profiles were calculated including sidechains (left) and only backbone atoms (right) respectively. Dashed lines show the profile for the starting structures and solid lines represent the simulation average with standard deviation in colored fill. The pore profiles for simulations of the with calcium X-ray structure and both cryo-EM structures overlap, and are in close agreement. The simulations started from the calcium free X-ray structure show a decrease in pore radius, largely attributable to backbone motion. Zoom in box (right) shows the most constricted region in the ECD with Trp407 and Asn405 being the two constricting residues.

**Figure S8:**
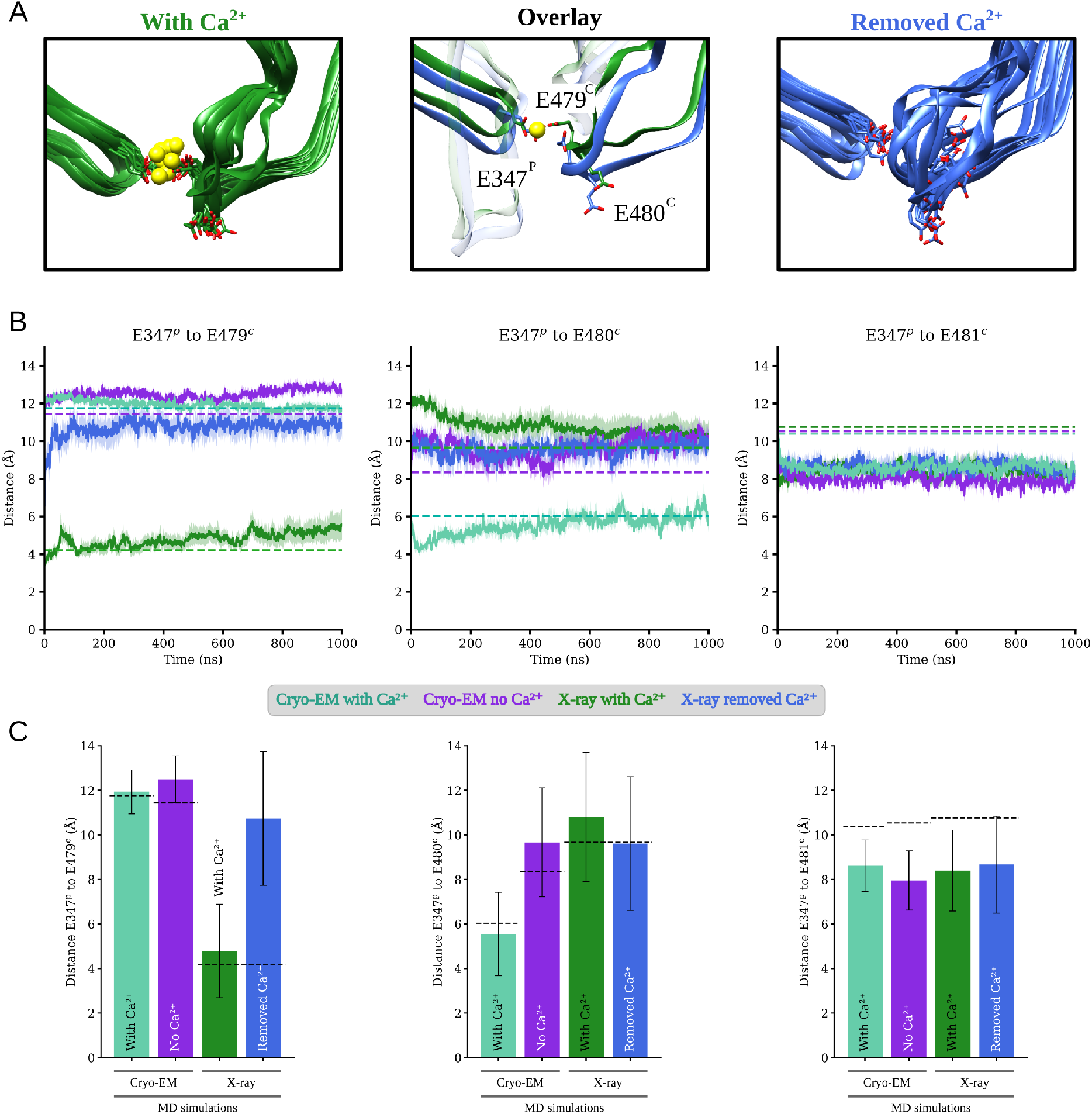
Behaviour of glutamates in the calcium binding site. (A) Snapshots from simulations starting from the with calcium X-ray structure by Hu et al. [11] (PDB ID: 6V4S); in green simulated with the resolved calcium ion present, and in blue simulated with the calcium ion removed. The center panel shows an overlay the initial simulation frame, the left and right panels show simulation snapshots spaced by 100 ns. (B) Distance between the sidechain carboxyl groups of glutamates in the calcium binding site, tracked through the MD simulations. The average value from the simulations, averaged over all subunit interfaces from four replicas, are shown as solid lines with the standard error of the mean in colorfill. Dashed lines show the values measured in the X-ray and cryo-EM structures. (C) Distance between the sidechain carboxyl groups of glutamates in the calcium binding site, averaged over time, and all subunit interfaces from four replicas. Error bars show standard deviation. Dashed lines show the distance in the cryo-EM or X-ray structure from which the simulation was started.

**Figure S9:**
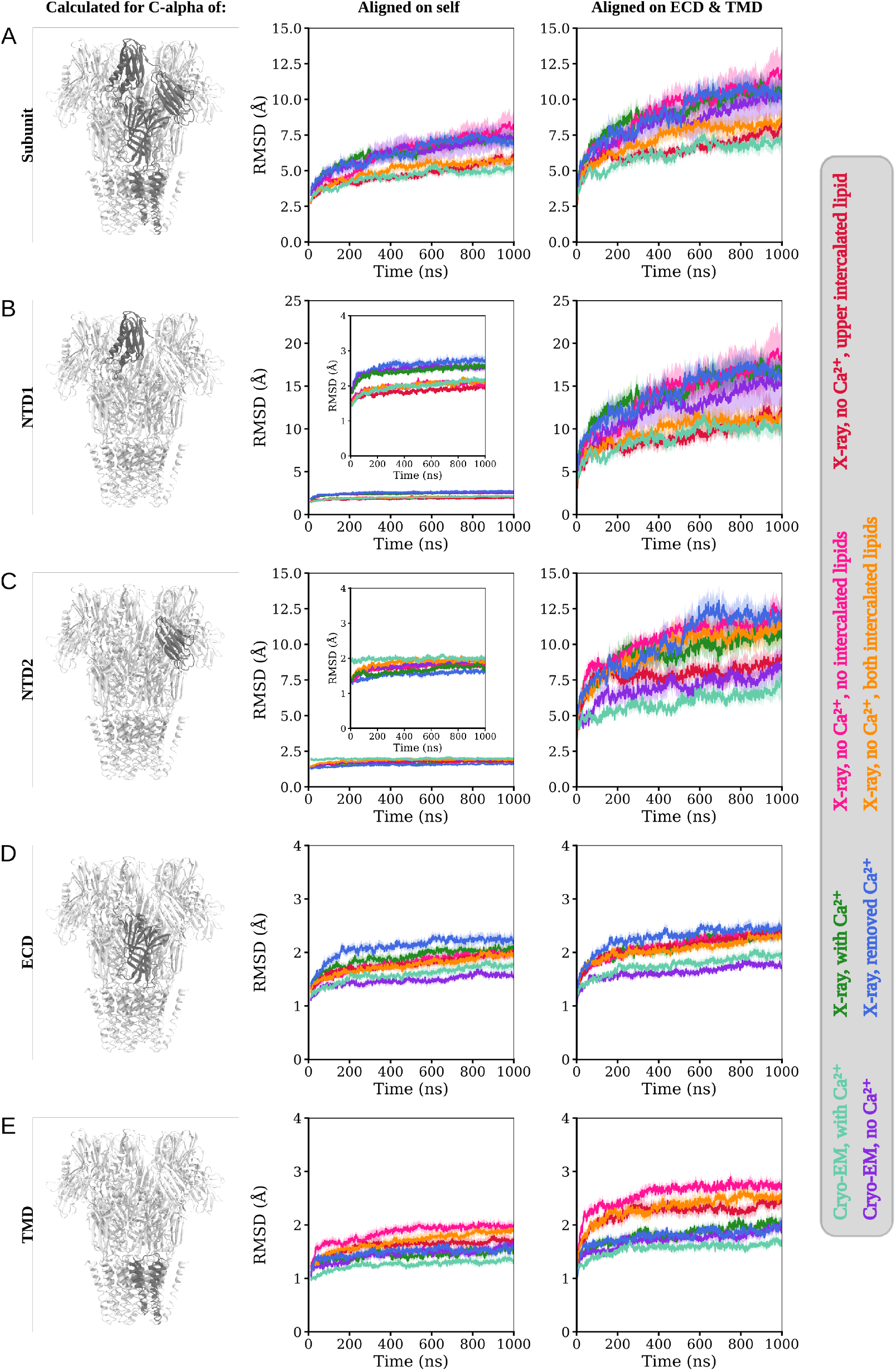
Root mean square deviation (RMSD) for the simulations of DeCLIC. Solid lines are averages over all subunits in four replicates, the standard error of the mean is shown as color fill. The RMSD has been calculated for the full subunit (row A), and for the four different domains in DeCLIC (rows B through E), aligning on either the domain itself (center column), or aligning on the joint ECD and TMD of the chain (right column). Rows are shown with the same scale for the RMSD values, inserts are shown in cases where the alignments yielded very different RMSD values. For the NTD lobes the RMSD is notably higher when aligning on the ECD-TMD compared to when alignment is done on the domain itself, indicating that these domains undergo translation relative to the ECD-TMD part of the protein.

**Figure S10:**
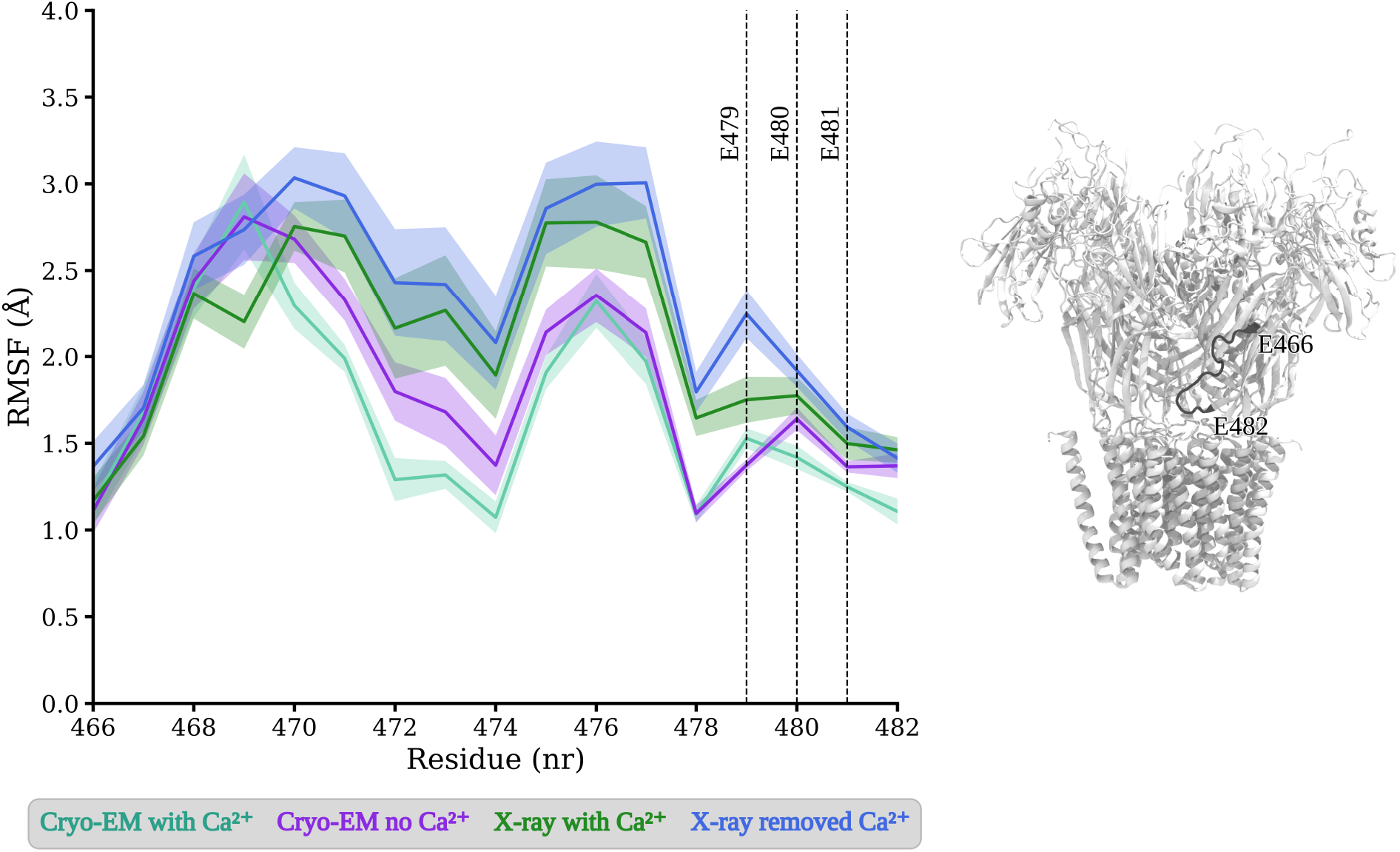
Root mean square fluctuation (RMSF) for residues in loop F. (residues shown in black on the structure shown to the right) from simulations starting from cryo-EM structures of DeCLIC in the presence (aquamarine) and absence (purple) of calcium, and from the X-ray structure with calcium - which was simulated with Ca^2+^ (green) and with Ca^2+^ removed (blue). Solid lines represent the average over all subunits in four replicas, and the standard error of the mean is shown as colorfill. Dashed vertical lines indicate the three consecutive glutamates E479, E480, and E481, which are located in the calcium binding site.

**Figure S11:**
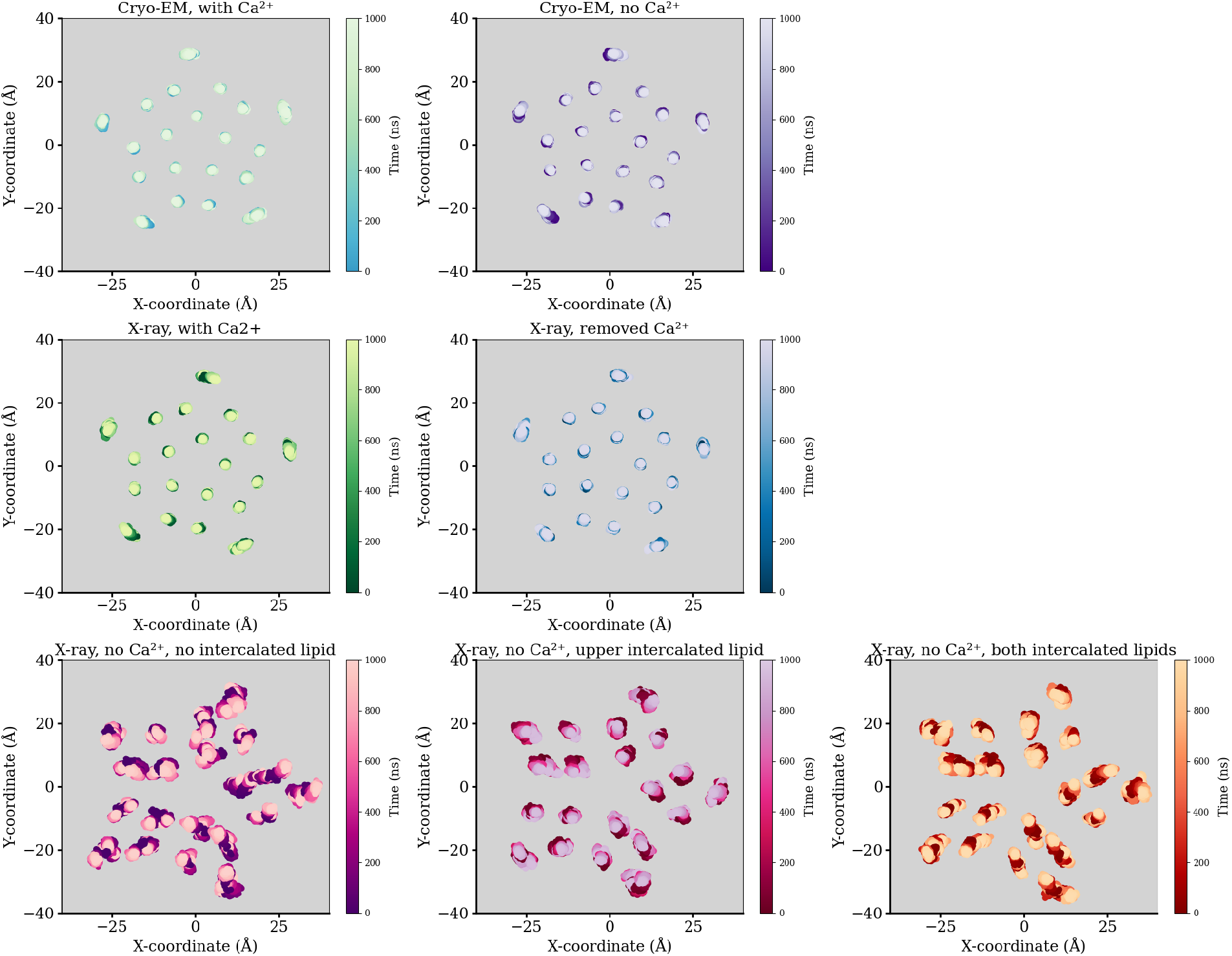
Center of mass position for the transmembrane helices through molecular dynamics simulations, four replicas, of the cryo-EM structures with and without Ca^2+^, the X-ray structure with Ca^2+^ (simulated with Ca^2+^ and with Ca^2+^ removed), and the X-ray structure without Ca^2+^ (simulated without intercalated lipid, with intercalated upper leaflet lipid, and with both upper and lower leaflet lipids intercalated). Simulations started from the cryo-EM structures and the with Ca^2+^ X-ray structure have stable helix positions through the simulations, with the most mobility for the lipid facing M4 helix. Simulations started from the no Ca^2+^ X-ray structure display less stable positions of the transmembrane helices.

**Figure S12:**
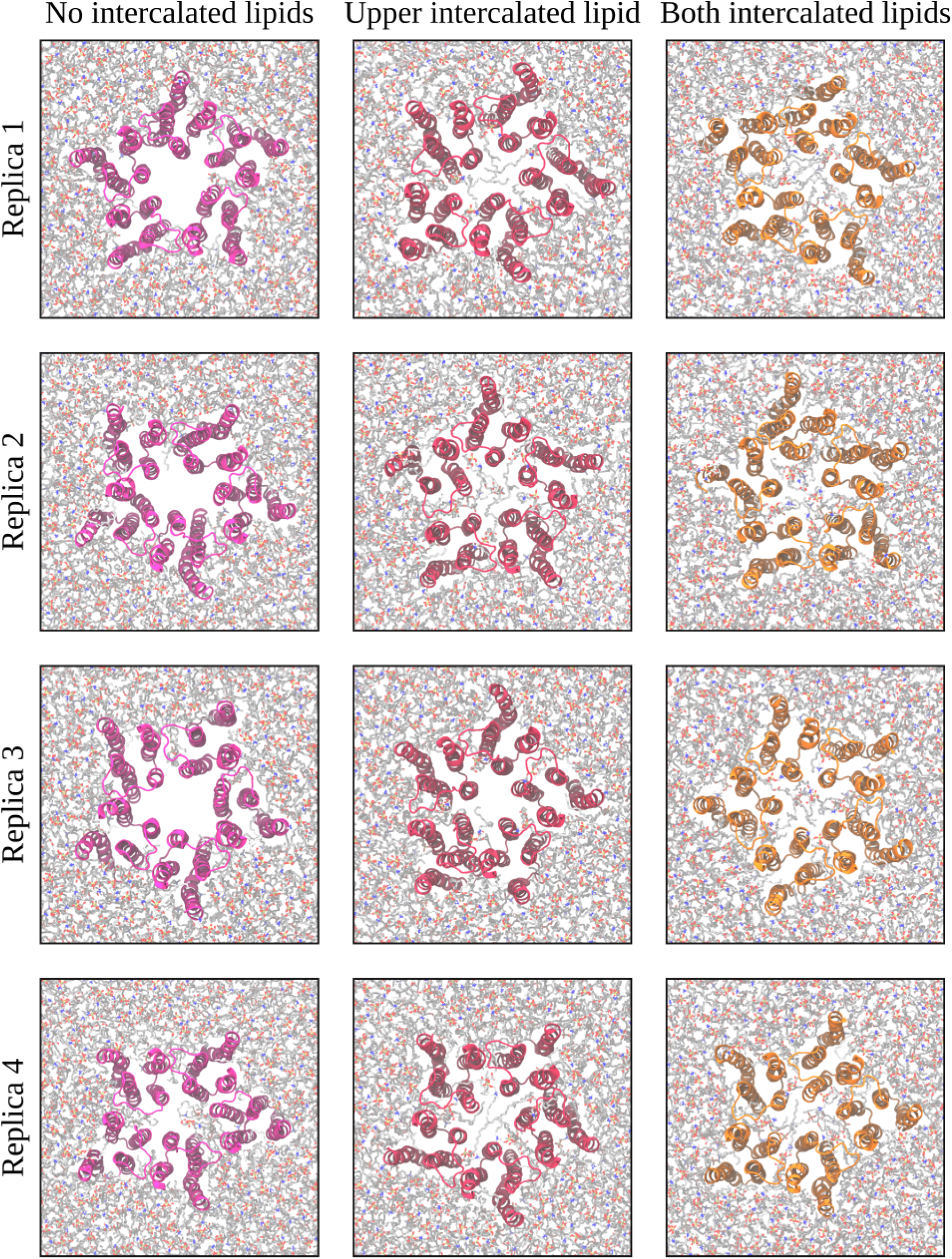
Final frame (1000 ns) of the simulations started from the X-ray structure with no calcium, showing the transmembrane domain and lipid bilayer seen from the extracellular side. The left column show simulations started without a lipid intercalated between the subunits, the center column simulations where an upper leaflet lipid had been intercalated under the M2-M3 loop, and the right column show simulations with both an upper and a lower leaflet lipid intercalated. The pore lining M2 helices deviated during the simulations from their symmetric starting positions, and especially the systems started with intercalated lipids were prone to have lipids penetrate into the pore.

**Figure S13:**
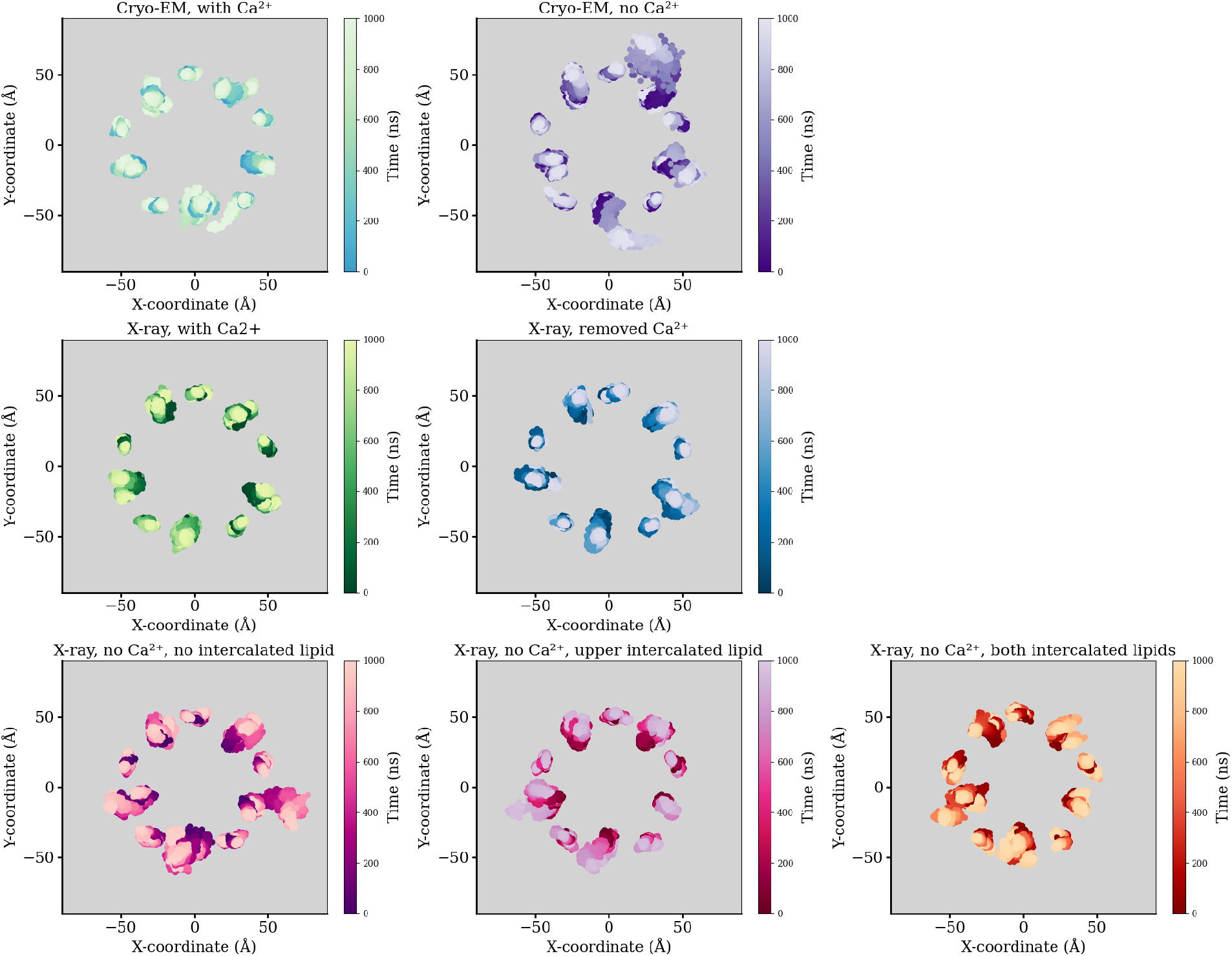
Position of the center of mass of NTD1 and NTD2 through the MD simulations, shown on the XY-plane. The larger clusters are from NTD1, and the smaller ones are from NTD2. Values from four replicate simulations are plotted for each system.

**Figure S14:**
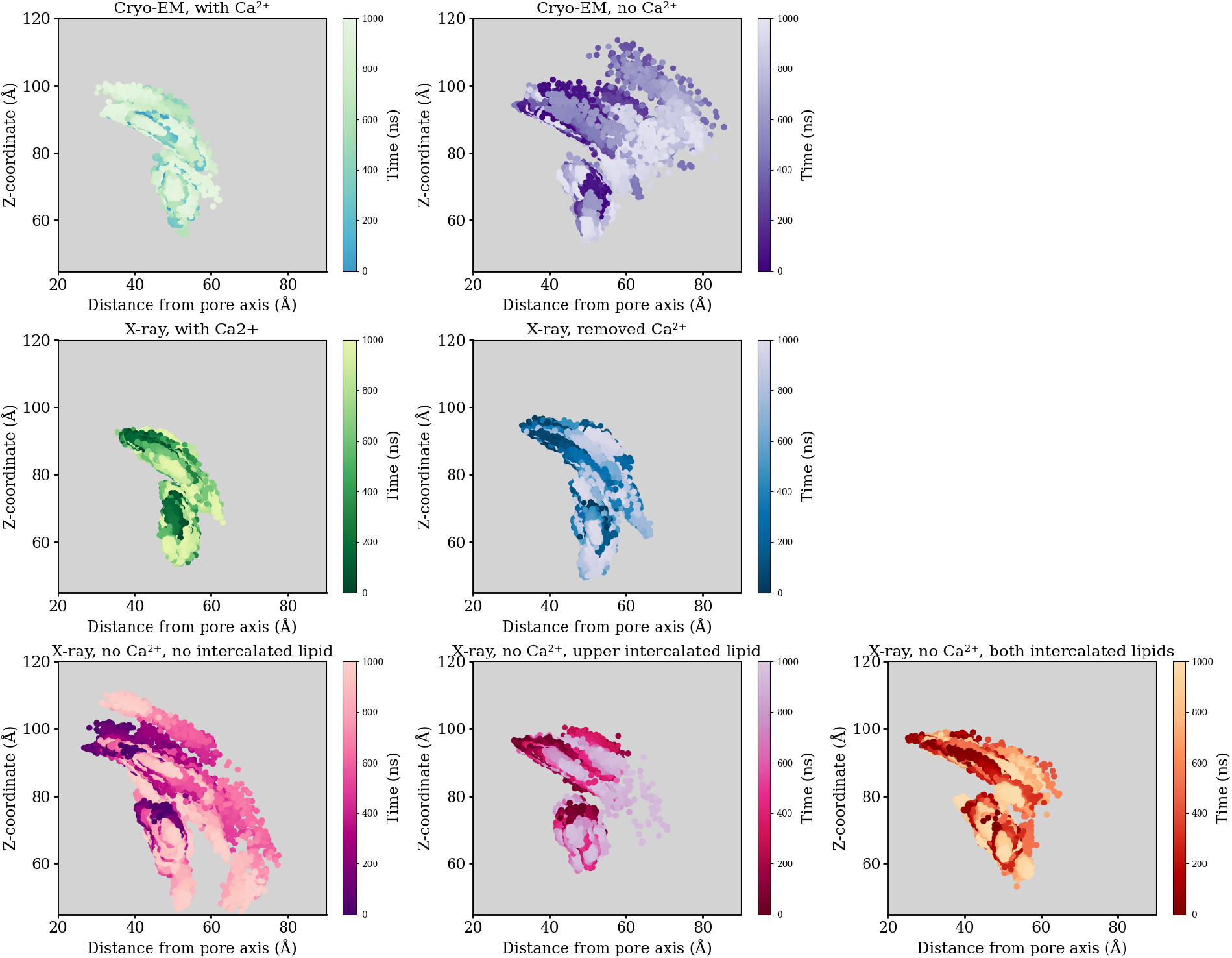
Position of the center of mass of NTD1 and NTD2 through the MD simulations, shown as Z-coordinate versus distance from pore axis. The upper cluster is from NTD1:s, and the lower from NTD2:s. Values from four replicate simulations are plotted for each system, and NTD lobes from separate subunits are individually represented.

**Figure S15:**
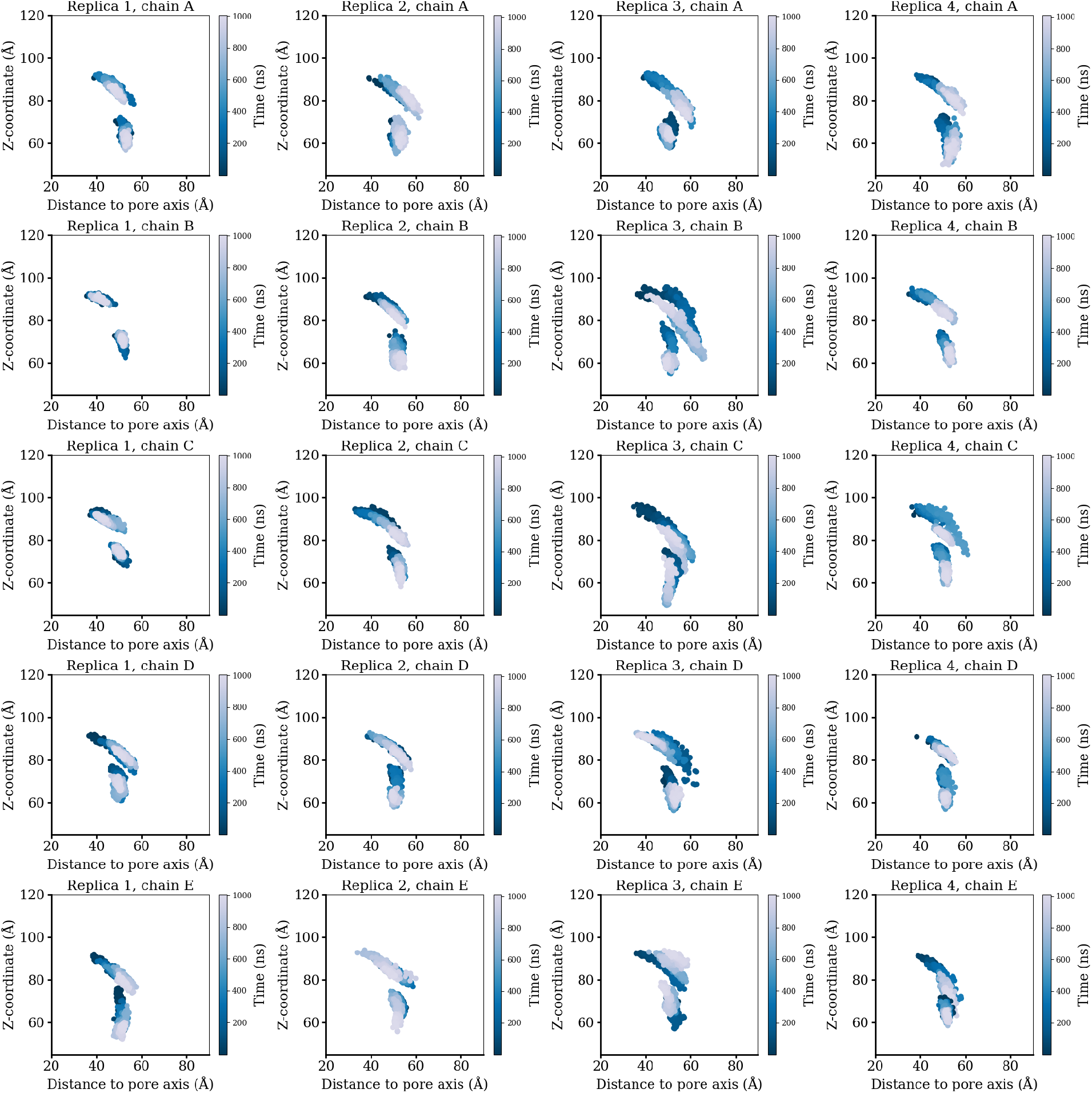
Center of mass position of NTD1 and NTD2 shown for individual subunits across four simulation replicas of the closed-like X-ray structure with Ca^2+^ removed for the simulations. The upper cluster is from NTD1, and the lower from NTD2. The simulations contain examples of the domains remaining close to the starting positions (e.g. replica 1 chain B: column 1, row 2), of NTD1 moving out-and-down and staying there (e.g. replica 2 chain C: column 2, row 3), and of NTD1 visiting the out-and-down conformation before returning to the starting position (e.g. replica 3 chain B: column 3, row 2).

**Figure S16:**
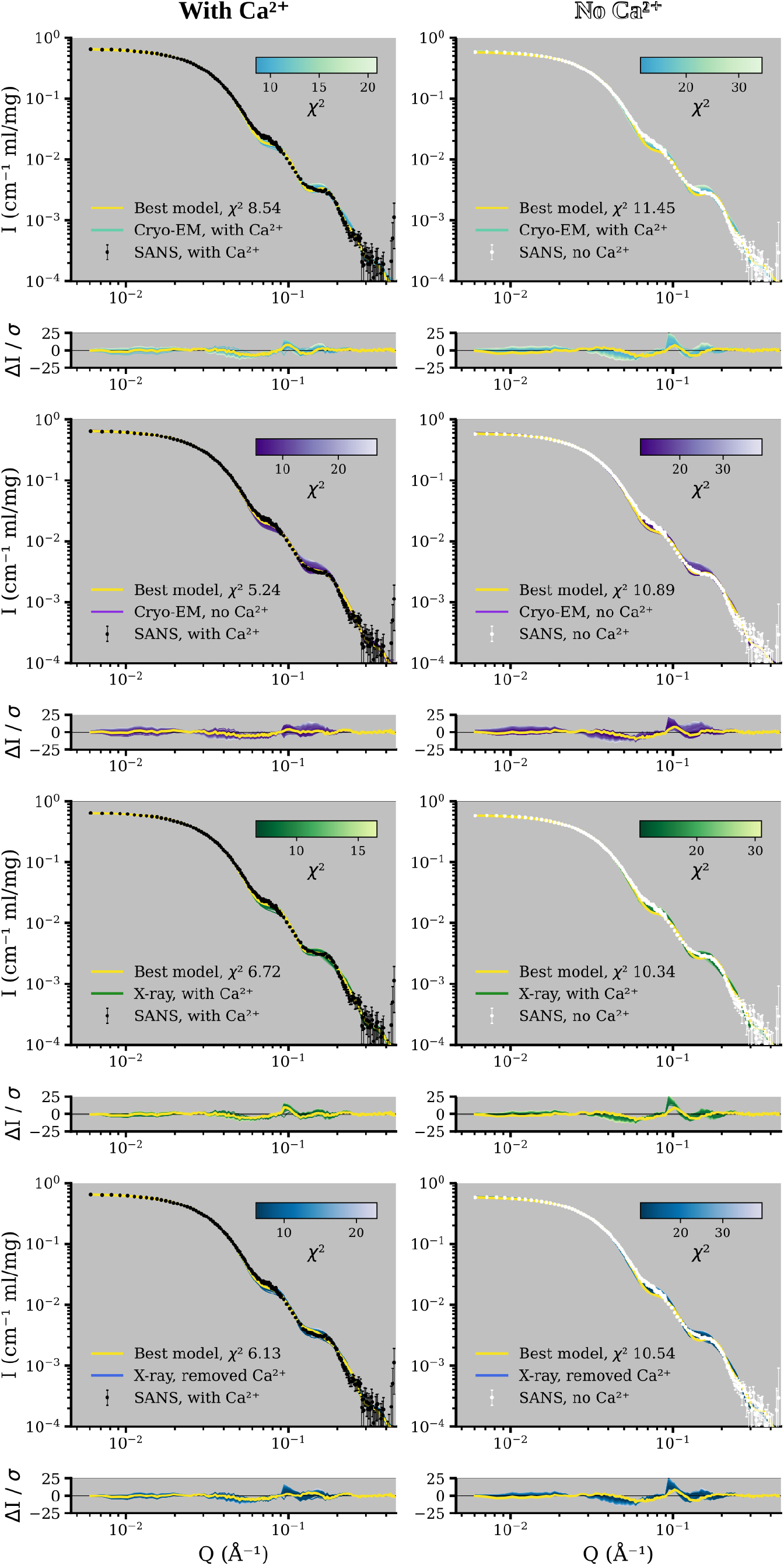
Fits to the SANS data collected with (black) and without (white) calcium for models from simulations started from closed-like structures, i.e. the cryoEM structures and the with calcium X-ray structure. For each simulation system the theoretical curve from the snapshot yielding the best goodness of fit to the SANS data is shown in yellow.

**Figure S17:**
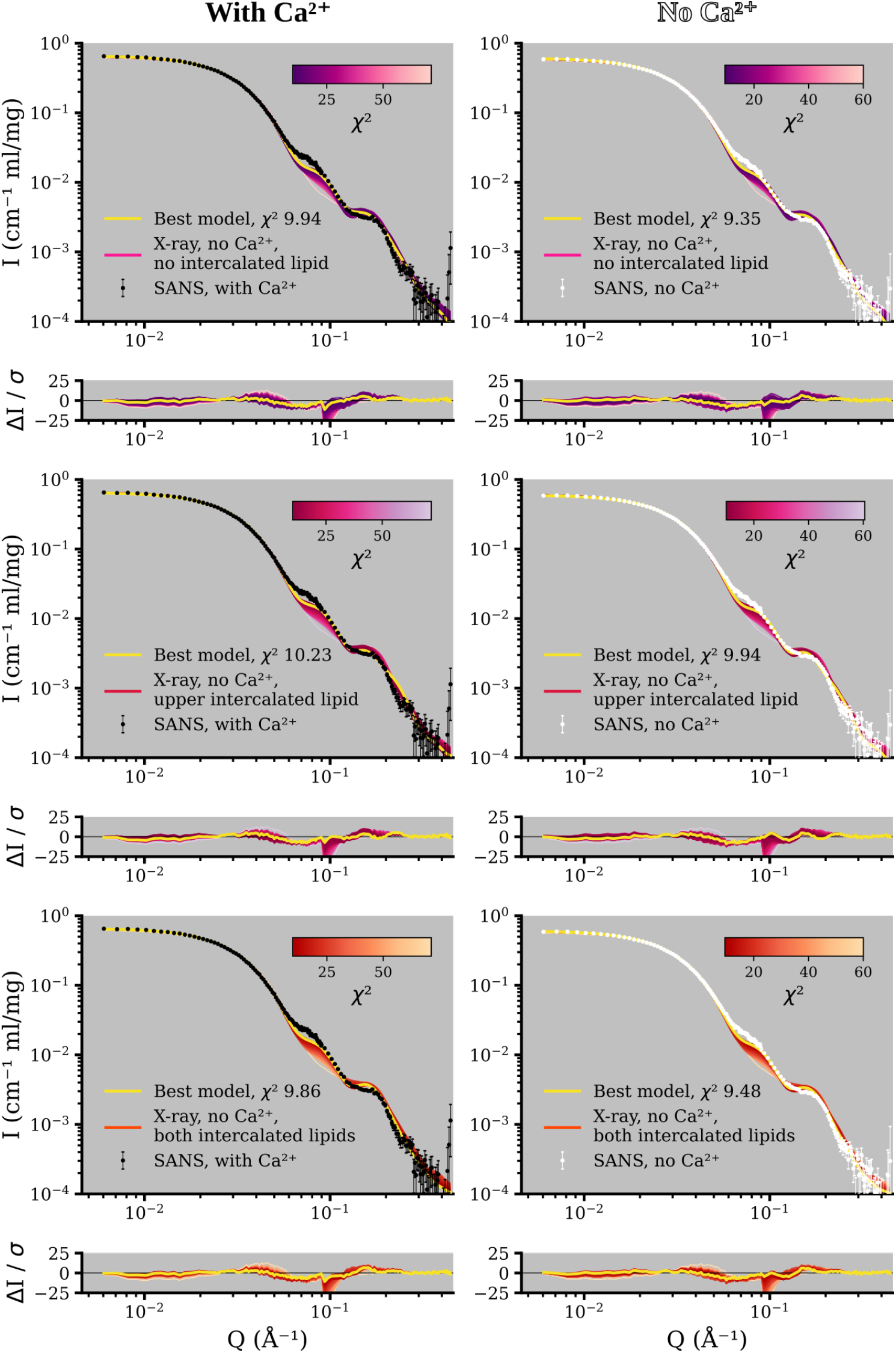
Fits to the SANS data collected with (black) and without (white) calcium for models from simulations started from the no calcium X-ray structure. For each simulation system the theoretical curve from the snapshot yielding the best goodness of fit to the SANS data is shown in yellow.

**Figure S18:**
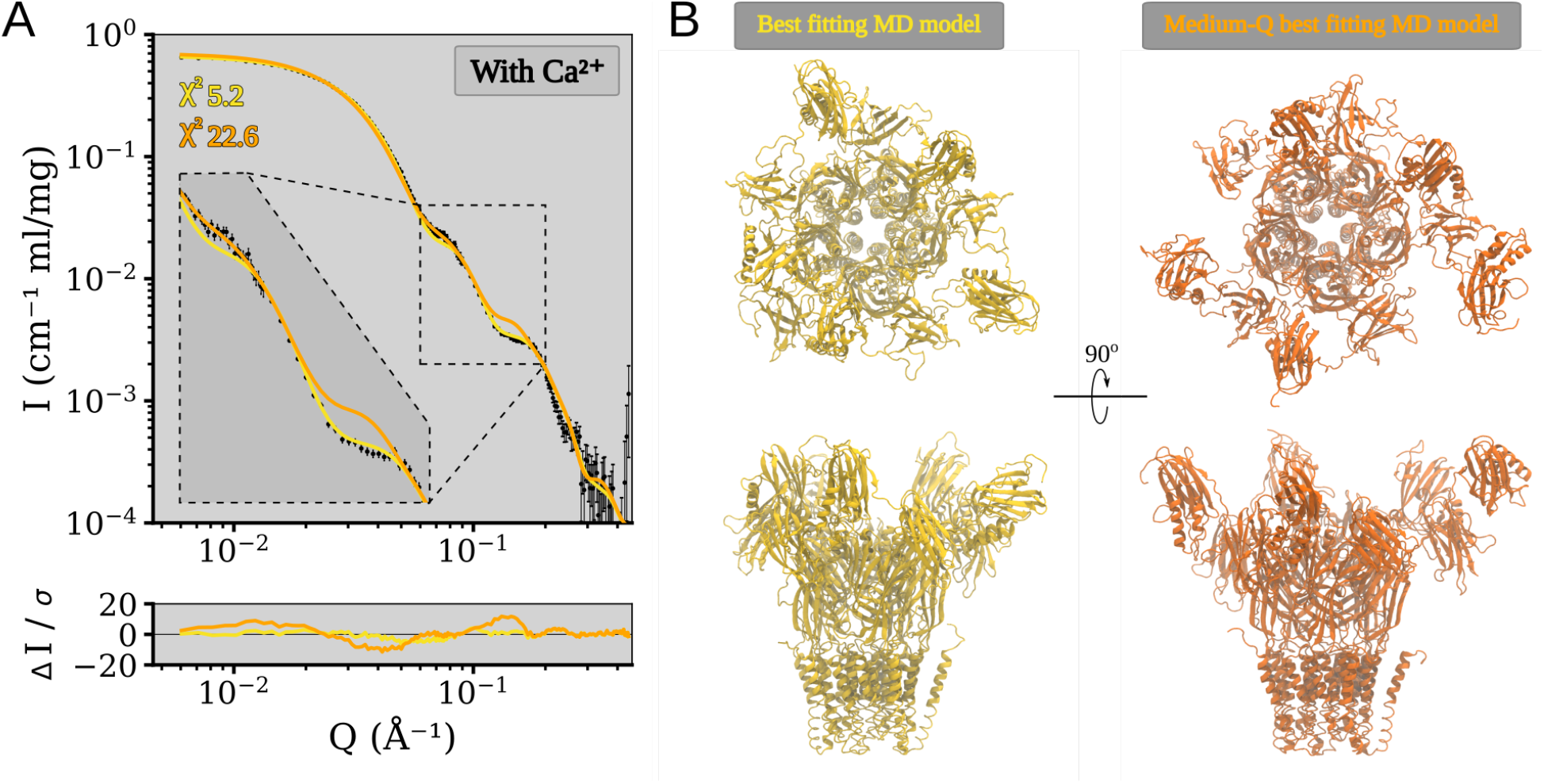
Comparison of the model with best overall fit to the with calcium SANS data and the model with the best fit to the *Q*∈ [0.06, 0.09] Å ^-1^ feature. (A) Theoretical curves from MD simulation snapshots of the cryoEM no Ca^2+^ structure, fitted to the with Ca^2+^ SANS data. In yellow is the snapshot which yielded the best over all fit to the SANS data (χ^2^ of 5.2), and in orange the one which had the best agreement with the medium-Q feature (*Q*∈ [0.06, 0.09] Å ^-1^, χ^2^ to full scattering profile of 22.6). The model with the best agreement with the medium Q-feature has worse fit at both higher and lower Q, as seen in the zoomed insert, and in the error weighted residual. (B) Top and side view of the the snapshot yielding the fits in panel A. Both models are asymmetric, and have different degrees of extension in the NTD.

**Movie S1:**
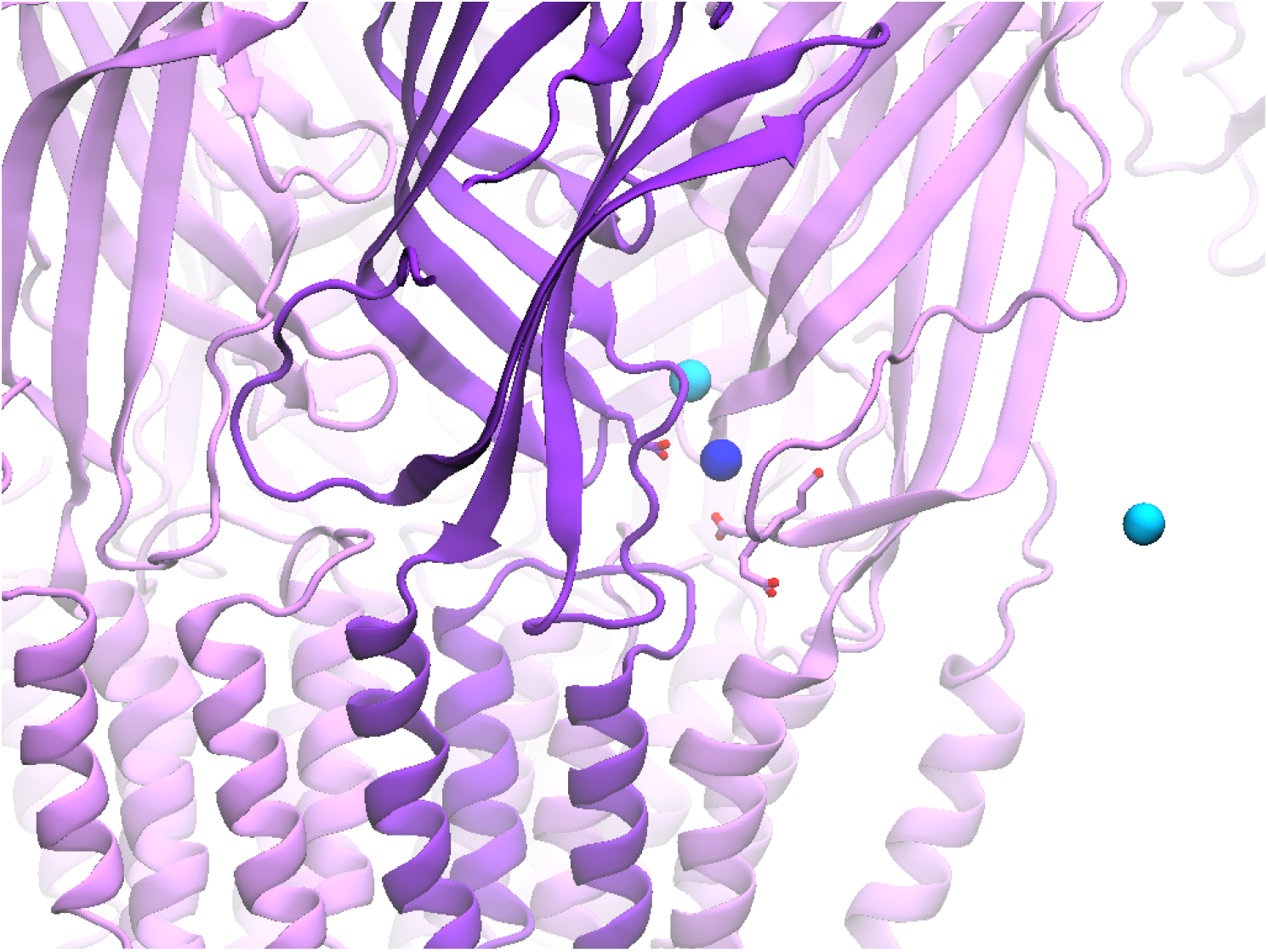
In the absence of calcium, sodium (cyan and blue) visit the calcium binding site, and some sodium ions pass between the central ECD vestibule and the outside of the protein through this site (one instance highlighted in blue). Glutamates in one calcium binding site are shown as sticks, and for clarity only sodium ions in the vicinity of this site are shown. The movie covers the final 300 ns of one replica, and was generated with trajectory smoothing on.

**Table S1:** Summary of structural parameters calculated from the SANS data. For the Guinier analysis the minimum and maximum Q-value included is also listed.

|  | With Ca<sup>2+</sup> | No Ca<sup>2+</sup> |
| --- | --- | --- |
| Guinier analysis |  |  |
| $I(0)$ (cm<sup>-1</sup>) | 0.68 | 0.62 |
| $R_g$ (Å) | 52.0 ± 0.2 | 51.6 ± 0.2 |
| $Q_{min}$ (Å<sup>-1</sup>) | 0.010 | 0.010 |
| $Q_{max}$ (Å<sup>-1</sup>), ($QR_g$ max) | 0.035 (1.7) | 0.035 (1.7) |
| Coefficient of correlation $R^2$ | 0.9997 | 0.9997 |
| $M$ from $I(0)$ (kDa), (ratio to predicted) | 348 (1.02) | 318 (0.93) |
| $P(r)$ analysis | | |
| $I(0)$ (cm<sup>-1</sup>) | 0.679 | 0.621 |
| $R_g$ (Å) | 51.26 ± 0.03 | 50.90 ± 0.03 |
| $d_{max}$ (Å) | 139.77 ± 0.74 | 139.26 ± 0.56 |
| $q$ range (Å<sup>-1</sup>) | 0.0060 - 0.4453 | 0.0060 - 0.4452 |
| $\chi^2$ | 0.91 | 0.71 |
| PepsiSANS |  |  |
| $R_g$ (Å) | 50.2 | 49.3 |

**Table S2:** Summary of modeling using protein structures and models based on protein structures, covering the radius of gyration of the model and the χ^2^ goodness of fit to the experimental data for the model.

| Structures | 6V4A | 6V4S | Cryo-EM<br>No Ca<sup>2+</sup> | Cryo-EM<br>With Ca<sup>2+</sup> |  |
| --- | --- | --- | --- | --- | --- |
| PepsiSANS |  |  |  |  |  |
| Predicted $R_g$ (Å) | 48.2 | 49.5 | 49.6 | 49.2 | |
| SANS with Ca<sup>2+</sup> $\chi^2$ | 79.23 | 10.75 | 11.18 | 12.12 | |
| SANS no Ca<sup>2+</sup> $\chi^2$ | 67.70 | 8.77 | 11.71 | 11.26 | |
| MD-simulation models |  |  |  |  |  |
|  | 6V4A<br>No Ca<sup>2+</sup> | 6V4S<br>With Ca<sup>2+</sup> | 6V4S<br>Removed Ca<sup>2+</sup> | Cryo-EM<br>No Ca<sup>2+</sup> | Cryo-EM<br>With Ca<sup>2+</sup> |
| Equilibration time (ns) | 2.25 | 2.25 | 2.25 | 2.25 | 2.25 |
| Simulation time (ns/replica) | 1000 | 1000 | 1000 | 1000 | 1000 |
| Replicas (nr) | 4 | 4 | 4 | 4 | 4 |
| $\chi^2$ range (PepsiSANS) | | | | | |
| SANS with Ca<sup>2+</sup> | 9.94 - 71.33 | 6.72 - 16.54 | 6.13 - 22.86 | 5.24 - 26.92 | 8.54 - 20.93 |
| SANS no Ca<sup>2+</sup> | 9.35 - 60.02 | 10.34 - 31.04 | 10.54 - 39.29 | 10.89 - 39.02 | 11.45 - 34.30 |

**Table S3:** Cryo-EM data collection and model refinement statistics

| *Data Collection* | **With Ca<sup>2+</sup>** | **No Ca<sup>2+</sup>** |
| --- | --- | --- |
| Microscope | FEI Titan Krios |  |
| Magnification | 165,000 |  |
| Voltage (kV) | 300 |  |
| Electron exposure (e<sup>-</sup>/Å<sup>2</sup>) | ~ 40 |  |
| Defocus Range (µm) | -2.0 – -3.6 | -1.4 – -3.2 |
| Pixel Size (Å) | 0.82 |  |
| Symmetry Imposed | C5 |  |
| Number of Images | ~ 2483 | ~ 11,505 |
| Particles Picked | ~ 156,000 | ~ 485,000 |
| Particles Refined | 19,731 | 43,650 |
| *Refinement* |  |  |
| Resolution (Å) | 3.5 | 3.2 |
| FSC Threshold | 0.143 |  |
| Map sharpening B-factor | -123 | -100 |
| Model Composition |  |  |
| Non-hydrogen Protein atoms | 19,570 | 21,620 |
| Protein Residues | 2585 | 2795 |
| Ligands | 5 | 0 |
| B-factor (Å<sup>2</sup>) | 33.4 | 67.5 |
| RMSD |  |  |
| Bond Lengths (Å) | 0.004 | 0.004 |
| Bond Angles (°) | 0.65 | 0.58 |
| Validation |  |  |
| Molprobity Score | 1.97 | 1.86 |
| Clashscore | 10.37 | 9.02 |
| Poor Rotamers (%) | 0 | 0 |
| Ramachandran Plot |  |  |
| Favored (%) | 93.4 | 94.4 |
| Allowed (%) | 6.6 | 5.6 |
| Outliers (%) | 0 | 0 |

**Table S4:** Table of missing residues and sidechains from the cryo-EM reconstructions

|  | With Ca<sup>2+</sup> | No Ca<sup>2+</sup> |
| --- | --- | --- |
| *Removed Residues* |  |  |
| NTD1 | 28 – 68 | 28 – 37 |
|  |  | 57 – 67 |
|  | 78 – 83 | 79 – 84 |
|  | 101 – 104 |  |
|  | 118 – 120 |  |
|  | 141 – 148 | 142 – 145 |
|  | 170 – 196 | 172 – 193 |
| NTD2 | 237 – 245 |  |
| C-terminus | 641 – 642 | 640 – 642 |
| *Total Removed* | **99** | **56** |
| *Removed Sidechains* |  |  |
| Leu | 75, 96, 100, 110, 197,<br>208, 257, 261, 271, 303 | 110, 197, 208, 235, 261,<br>311 |
| Ile | 86, 107, 117, 137, 151,<br>199, 279 | 137, 199 |
| Asp | 87, 105, 115, 122, 215,<br>263, 278, 290, 294, 343,<br>361, 418, 435, 437 | 76, 115, 140, 213, 276,<br>361 |
| Glu | 97, 134, 209, 270, 297,<br>313, 425 | 46, 241, 270, 313 |
| Arg | 116, 211, 224, 301, 312 | 211 |
| Lys | 127 | 98, 127, 158, 214 |
| Ser | 132 | 132 |
| Phe | 135, 259 | 259 |
| Asn | 153, 310 | 104 |
| Gln | 277, 316 | 316 |
| Val | 319, 323 | 319 |
| Thr |  | 71 |

**Table S5:** Sample details.

|  |  |
| --- | --- |
| Organism of origin | *Desulfofustis* sp. PB-SRB1 |
| Expression system | *Echerichia coli* |
| UniProt ID | V4JF97 |
| Extinction coefficient [$A_{280}$ 0.1%(w/v)] | 1.107 |
| Volume from structure ($\text{\AA}^3$) | 495062 |
| Particle contrast from sequence and solvent constituents, $\Delta\bar{\rho}$ ($\vert \rho_{protein} - \rho_{solvent}\vert $; $10^{10} \text{ cm}^{-2}$) | 3.94<br>($\vert 2.44 - 6.38\vert $) |
| M from chemical composition (kDa) | 341.7 |

**Table S6:** Summary the SANS collection parameters, including wavelength, pathlength, detector distances, Q-range, absolute scaling method, normalization, sample concentration and volume, and buffer composition.

|  |  |
| --- | --- |
| Instrument | ILL D22 |
| Wavelength ($\text{\AA}$) | 6 |
| Pathlength (cm) | 0.1 |
| Detector distances (m) | 2.8m/2.8m & 8m/8m |
| $Q$ measurement range ($\text{\AA}^{-1}$) | 0.006 - 0.452 |
| Absolute scaling method | Incident beam flux |
| Normalization | Divided by concentration |
| Exposure time (aggregate time) |  |
| With $\text{Ca}^{2+}$ | 111 x 30s ( 55.5 min) |
| No $\text{Ca}^{2+}$ | 127 x 30s (63.5 min) |
| Column | Superdex 200 Increase 10/300 |
| Sample temperature ($^{\circ}\text{C}$) | 10 |
| Loading concentration | 6 mg/ml |
| Injection volume | 240 $\mu\text{l}$ |
| Flow rate (ml/min) | 0.2, 0.01, 0 |
| Average concentration (mg/ml) in combined data frames |  |
| With $\text{Ca}^{2+}$ | 1.09 |
| No $\text{Ca}^{2+}$ | 1.14 |
| Solvent |  |
| With $\text{Ca}^{2+}$ | $\text{D}_2\text{O}$, 150 mM NaCl, 20 mM Tris·HCl, 10 mM $\text{CaCl}_2$, 0.5 mM d-DDM |
| No $\text{Ca}^{2+}$ | $\text{D}_2\text{O}$, 150 mM NaCl, 20 mM Tris·HCl, 10 mM EDTA, 0.5 mM d-DDM |

**Table S7:**
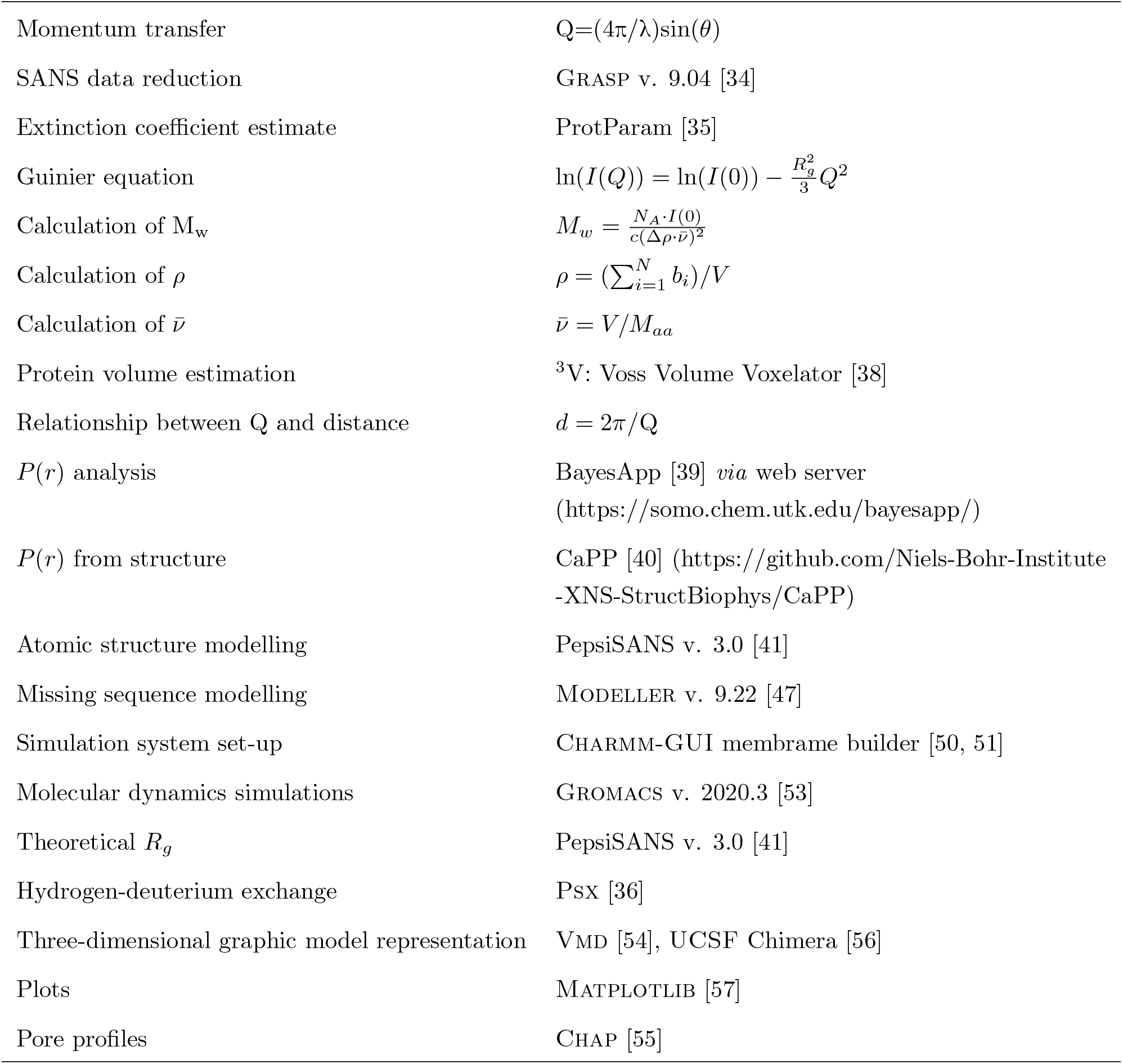
Summary of software and equations employed for SANS data reduction, analysis, and interpretation.

## Notes

https://doi.org/10.5281/zenodo.6022369

https://www.sasbdb.org/project/1572/

https://doi.ill.fr/10.5291/ILL-DATA.8-03-1002

### Competing Interest Statement

The authors have declared no competing interest.

